# Monosomy X in isogenic human iPSC-derived trophoblast model impacts expression modules preserved in human placenta

**DOI:** 10.1101/2021.12.13.472325

**Authors:** Darcy T. Ahern, Prakhar Bansal, Isaac Faustino, Yuvabharath Kondaveeti, Heather R. Glatt-Deeley, Erin C. Banda, Stefan F. Pinter

## Abstract

Mammalian sex chromosomes encode homologous X/Y gene pairs that were retained on the male Y and escape X chromosome inactivation (XCI) in females. Inferred to reflect X/Y-pair dosage sensitivity, monosomy X is a leading cause of miscarriage in humans with near full penetrance. This phenotype is shared with many other mammals but not the mouse, which offers sophisticated genetic tools to generate sex chromosomal aneuploidy but also tolerates its developmental impact. To address this critical gap, we generated X-monosomic human induced pluripotent stem cells (hiPSCs) alongside otherwise isogenic euploid controls from male and female mosaic samples. Phased genomic variants of these hiPSC panels enable systematic investigation of X/Y dosage-sensitive features using *in vitro* models of human development.

Here, we demonstrate the utility of these validated hiPSC lines to test how X/Y-linked gene dosage impacts a widely-used model for the human syncytiotrophoblast. While these isogenic panels trigger a *GATA2/3* and *TFAP2A/C* -driven trophoblast gene circuit irrespective of karyotype, differential expression implicates monosomy X in altered levels of placental genes, and in secretion of placental growth factor (PlGF) and human chorionic gonadotropin (hCG). Remarkably, weighted gene co-expression network modules that significantly reflect these changes are also preserved in first-trimester chorionic villi and term placenta. Our results suggest monosomy X may skew trophoblast cell type composition, and that the pseudoautosomal region likely plays a key role in these changes, which may facilitate prioritization of haploinsufficient drivers of 45,X extra-embryonic phenotypes.

## INTRODUCTION

Placental female mammals maintain dosage parity of most X-linked genes with males via X chromosome inactivation (XCI), a process which independently evolved long non-coding RNAs (eutherian *XIST*, metatherian *Rsx*) to silence one of two X chromosomes (1). This dosage-compensation strategy was likely necessitated by attrition of the proto-Y in the heterogametic male germline of ancestral mammals following acquisition of the male-determining factor (*SRY*) (2). Yet, because some sex-chromosomal genes were selected to resist Y-attrition and escape XCI, thereby maintaining expression from two active copies in males and females alike (3), mammalian development is likely sensitive to proper dosage of these “X/Y-pair” genes (4).

This multi-genic dosage-sensitivity is perhaps best reflected in the pronounced rarity of live-born monosomy X in mammals that feature a long pseudo-autosomal region (PAR). Present on X and Y, the PAR is maintained via meiotic recombination in the male germline, and likewise escapes XCI in females (5). In contrast, mice feature a short PAR, as well as comparatively few XCI “escapee” genes overall (6, 7), and tolerate monosomy X with very little developmental impact (8, 9).

Human monosomy X (45,X) causes Turner syndrome (TS, ~1:2500 live births), which ranges from full penetrance of short stature and early, often pre-pubertal ovarian failure, to other skeletal and craniofacial changes, lymphedema of hands and feet, cardiovascular defects, and impaired hearing in about half of TS patients (10, 11). Yet, most monosomy X pregnancies result in miscarriage, estimated to account for 6-11% of all spontaneous terminations (12–15). Because the rate of detectable mosaicism for euploid cells in live-born TS is very high (~50%), but very low (~0.5%) in karyotypic follow-up of miscarried monosomy X, Hook and Warburton compellingly hypothesized zygotic monosomy X to be near-invariably lethal *in utero*, and that live-born TS results from mitotic sex chromosome loss in early embryos, which gave rise to either detectably mosaic TS, or TS with cryptic (e.g. confined placental) mosaicism (15).

Conversely, this would suggest that it in absence of placental mosaicism, homogenously 45,X extra-embryonic tissues would be defective in supporting conceptuses to term. Because this phenotype is not shared with the mouse, new mammalian and human *in vitro* models are needed to address this important question. While two prior reports of 45,X human embryonic and induced pluripotent stem cell (hiPSC) lines pointed to lower expression of some placental genes in non-directed (embryoid body) differentiation (16, 17), the impact of monosomy X on relevant *in vitro* models of human extra-embryonic development has not been assessed.

To address this important question in a widely-used hiPSC-based model of the primitive syncytiotrophoblast (STB) (18, 19), we derived 45,X hiPSCs alongside isogenic euploid control lines from mosaic samples. This enables us to largely exclude the impact of autosomal variation, and leverage phased genome sequencing to quantify allele-specific dosage contributions from X and Y. While these hiPSCs trigger a *GATA2/3, TFAP2A/C*-mediated gene circuit (20) irrespective of karyotype, differentially-expressed genes indicate monosomy-X alters the balance between cytotrophoblast (CTB), STB and extravillous trophoblast (EVT) markers. These changes are also reflected in secretion of human chorionic gonadotropin (hCG) and placental growth factor (PlGF), and correlated gene modules are preserved in late first-trimester and term placentas. Together, our study represents the largest single source of 45,X hiPSC and isogenic euploid control lines to-date, and provides a first direct assessment of how monosomy X may impact human trophoblast-relevant gene networks.

## RESULTS

Because the low rate of term pregnancy with 45,X conceptuses may reflect a selective bottleneck, we turned to a post-developmental model of sex chromosome loss, namely aging, as a source of mosaic human monosomy X, which by age 75 increases to ~45% and ~0.45% in males and females, respectively (21–23). In total, we reprogrammed mosaic fibroblasts from four donors (three female, one male) and validated the resulting clones in systematic fashion (Fig. 1A). Two of the female samples resulted in exclusively X-monosomic or euploid hiPSC lines, as determined by tandem repeat PCR or copy-number quantitative PCR (qPCR) at X-linked loci (data not shown). These lines were not pursued further, as our analysis aimed to exclude effects of autosomal genetic variation by comparing 45,X to matched isogenic euploid control lines of the same donor. In contrast, female (AG05278) and male (AG09270) samples resulted in both euploid and 45,X hiPSC clones (XO, XO2/8/9) from each donor (aged 65 & 67). Karyotyping (Fig. S1A) further ruled out three male-derived clones with chromosome (chr) 12 trisomy or duplications. Finally, all remaining male and female-derived lines passed high-resolution cytogenetic testing (CytoSNP-850k) to rule out any other genomic copy-number variation (CNV). No other CNV calls (at minimal CNV resolution of 400 kb) were made in AG09270-derived lines, and a single 440 kb duplication in neuron-specific *OPCML* was found in all AG05278-derived hiPSCs lines and donor fibroblasts, yielding a final set of eight hiPSC clones with validated karyotypes (Fig. 1B).

**Figure 1:**
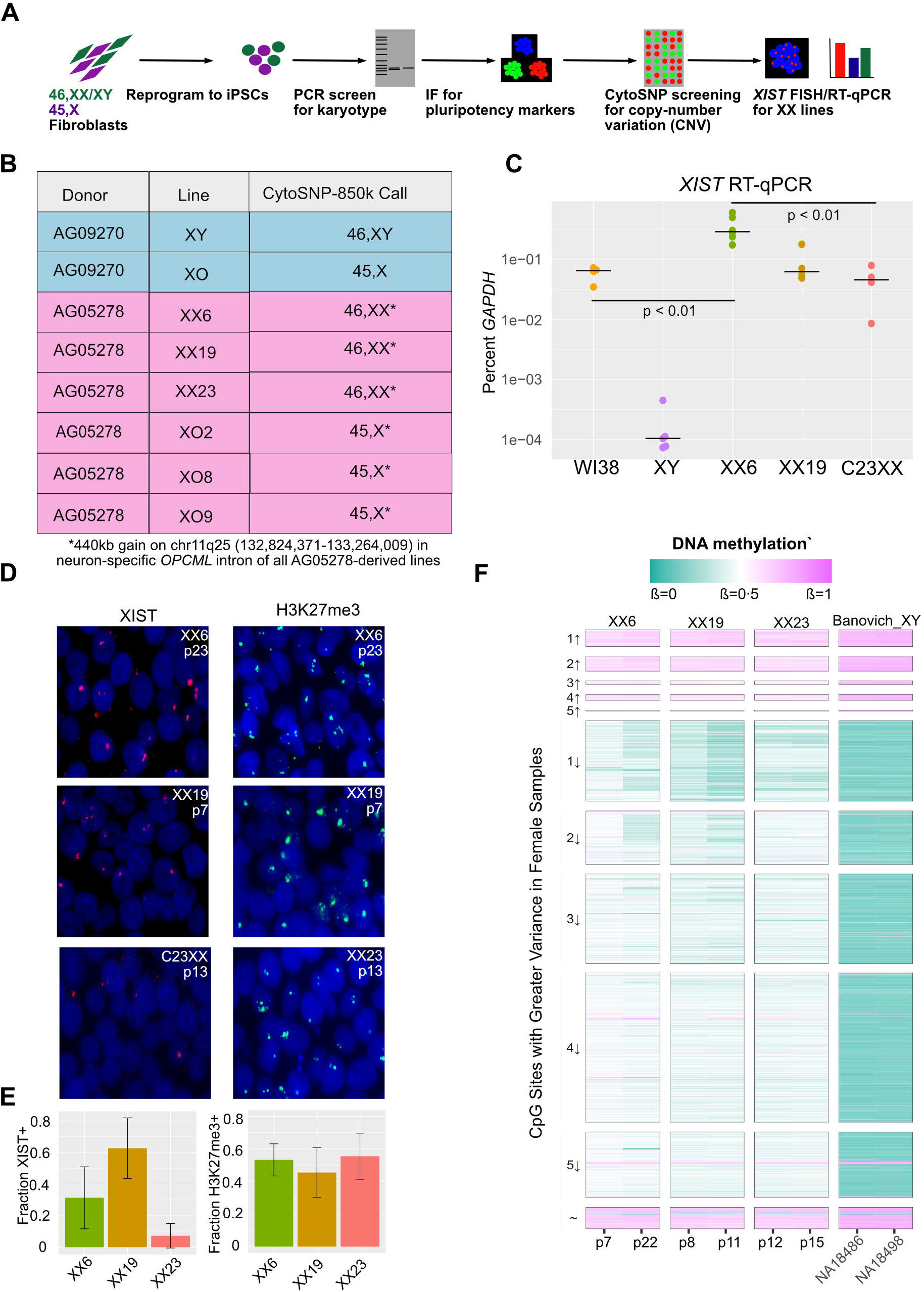
**A.)** Schematic of reprogramming and hiPSC characterization. **B.)** Cytogenetic characterization (CytoSNP-850k, Illumina) of hiPSC clones used in this study. **C.)** *XIST* expression by RT-qPCR as a percentage of *GAPDH* (Mann-Whitney-U test p-values). **D.)** *XIST* FISH (left) and H3K27me3 immunofluorescence (IF, right) images of three 46,XX hiPSC lines. **E.)** Quantification of *XIST+* (left) and H3K27me3+ (right) cells. **F.)** X chromosome DNA methylation levels across all three XX clones at two passages inclusive to all experiments reported herein, as well as XY hiPSC data from (118). Depicted probes indicate methylated DNA fraction (β value) labeled by associated transitions (1 through 5) and change in DNA methylation during Xi erosion (from (26)).

We next characterized XCI in female-derived 46,XX clones (XX6, XX19, XX23), to confirm intact X dosage compensation. Loss of *XIST* expression is common in standard hiPSC culture, and can lead to progressive reactivation of the inactive X (Xi), referred to as Xi erosion (24, 25). We recently reported a prevalent and contiguous Xi DNA hypomethylation trajectory after loss of *XIST* expression, which we validated in two of our 46,XX (XX19/23) lines over six months of continuous passage (26). However, in early-passage and differentiation experiments described herein, all three 46,XX clones expressed *XIST* at or above female fibroblast WI-38-equivalent levels by qPCR (Fig. 1C), and by fluorescence *in-situ* hybridization (FISH) (Fig. 1D). As expected based on our and prior reports, these early passages reflect heterogeneity for *XIST* levels and associated H3K27me3-deposition on the Xi (Fig. 1E). We therefore confirmed that X-linked CpGs that best reflect Xi erosion (26) remain largely hypermethylated in all three hiPSCs lines (Infinium MethylationEPIC), indicating intact XCI maintenance inclusive to all early hiPSC passages (XX6 < p21, XX19 < p9, XX23 < p11) used in experiments described herein (Fig 1F). To enable allele-specific assessment of XCI across individual samples in subsequent RNA-seq experiments, we also performed linked-read whole-genome sequencing on the 10X Genomics platform (phased WGS to ~30x coverage), yielding a catalogue of 76,737 heterozygous variants that distinguish the X chromosomes in female-derived lines (Fig. S1B). In the mRNA-seq experiments below, we leverage these phased variants (1,056 in exons, 3’ and 5’ UTRs) to determine that all female 45,X clones maintained the same parental X chromosome, and to quantify escape from XCI and X reactivation.

We confirmed high expression of pluripotency markers by immunofluorescence (IF) staining for OCT4 and SSEA4, and low expression of markers associated with germ layer differentiation by 3’ mRNA sequencing (Fig. S2A,B). All 45,X hiPSC lines expressed high levels of pluripotency genes *SOX2* and *DNMT3B* and low levels of lineage-specific genes, equivalent to their respective euploid control lines, and in line with the few 45,X lines described previously (17, 27–29). Several methods for derivation of trophoblast-like (TBL) cell fates from hiPSCs have been reported in recent years (reviewed in (30)), starting from either pre-implantation blastocyst-like (naïve or ground state) (31, 32), post-implantation epiblast-like (or primed) (18, 33, 34), or intermediate (extended/expanded) (35, 36) human pluripotency states. We chose a well-characterized TBL-induction method compatible with primed hiPSCs (18, 33), to ensure our validated karyotypes and DNA methylation (Fig. 1) were stably maintained. These requirements ruled out TBL-induction from naïve hiPSCs, which can suffer increased genome instability relative to primed hiPSCs (37, 38), and genome-wide loss of DNA methylation (39, 40) that includes imprinted genes (41), to which extra-embryonic development is highly dosage-sensitive (42–45).

Primed hiPSCs were differentiated to TBL cell fates by exposure to BMP4 and inhibition of TGFb1/activin/nodal signaling over 8 days. This widely used “BAP” (BMP4, A83-01, PD173074) model (18, 33, 34, 36, 46–48) is thought to reflect the primitive STB (18, 48), as BMP4 activates a conserved trophectodermal transcription factor (TF) circuit via *GATA2/3* and *TFAP2A/C* (20, 49, 50). All BAP-treated lines formed flat epithelial sheets that gave rise to mononuclear HLA-G^+^ cells (Fig. S3A), or progressively fused to form large syncytiated cells, seen as clustered nuclei inside Na+/K+ ATPase-marked membranes (Fig. 2A). These syncytia expressed high levels of the hCG beta subunit, which is only produced by the fused STB upon implantation and is important for endometrial receptivity and maternal immune suppression (51, 52). Abnormal hCG concentrations have been associated with adverse pregnancy outcomes, including intrauterine demise and fetal growth restriction (IUGR), as well as pre-eclampsia (53, 54). All BAP-treated cells secreted high levels of hCG, with a moderate but statistically significant decrease in 45,X lines (XO, XO2/8/9) relative to their isogenic euploid controls by ELISA (Fig. 2B). Another important protein secreted from the STB, placental growth factor (PlGF) is a member of the vascular endothelial growth factor (VEGF) family, and known to be critical for proper placental angiogenesis (55). PlGF levels were also lower in 45,X relative to their isogenic euploid controls, a decrease that was statistically significant in the female panel (XX19/23 euploid vs. XO2/8/9), and in comparing all 45,X to XX19/23 and XY samples (Fig. 2C, Mann-Whitney-U p = 9.2e-05).

**Figure 2:**
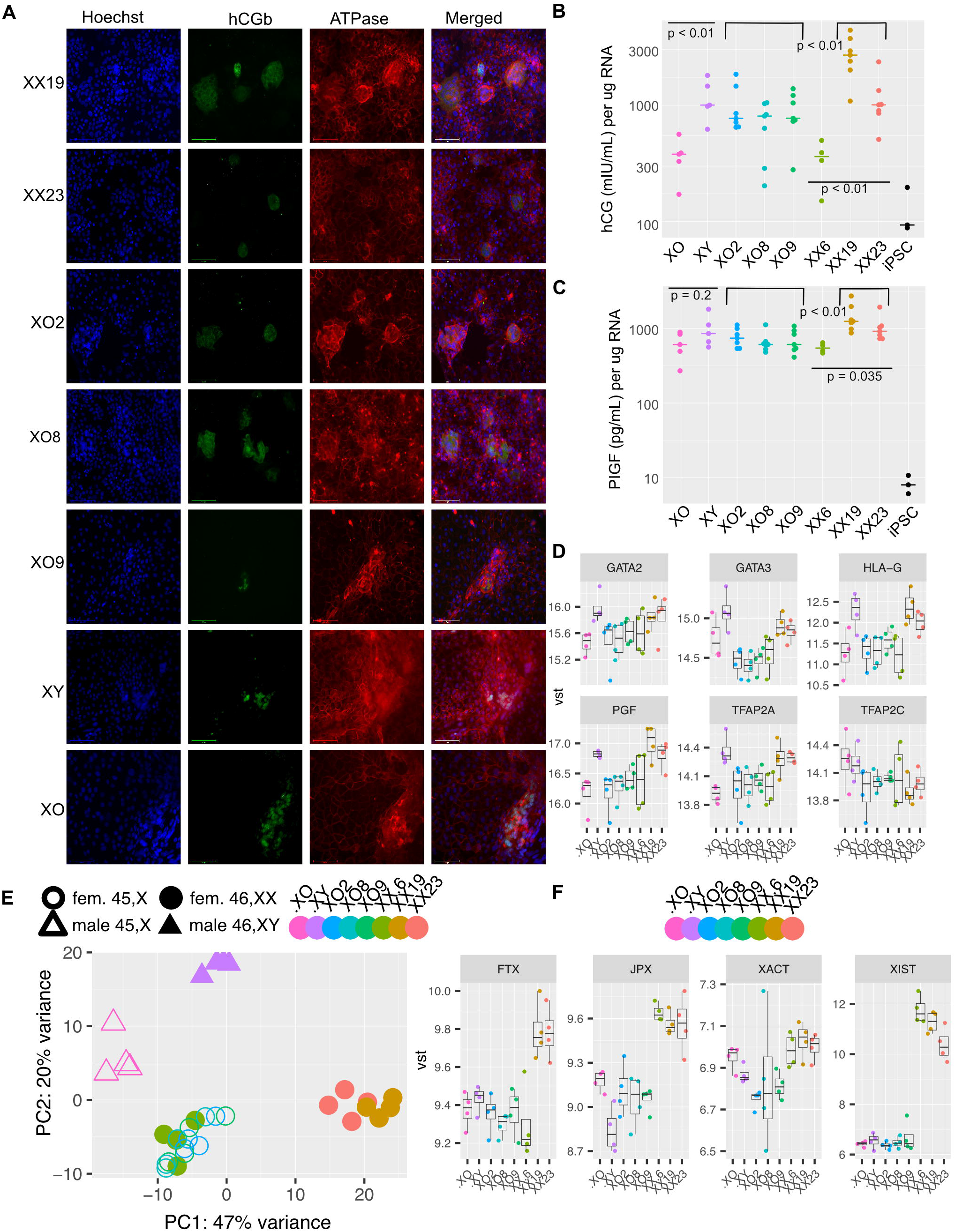
**A.)** IF images of TBLs stained for hCGb and membrane-marking Na+/K+ ATPase, nuclei counterstained with Hoechst. **B.)** hCG ELISA results in mIU/mL media per ug of RNA harvested from the same well. **C.)** PlGF ELISA results in pg/mL per ug of RNA, as in B. Mann-Whitney-U tests p-values reported in B and C compare 45,X samples to otherwise isogenic euploid controls (as denoted by brackets). **D.)** TBL RNA-seq vst counts of genes relevant to BAP-induced cell fates, plotted by line. **E.)** Principal component analysis (PCA) of 45,X and otherwise isogenic euploid control TBL RNA-seq data. Respective symbols and colors indicate karyotype and cell line. **F.)** TBL RNA-seq vst counts of X-linked non-coding RNA genes relevant to XCI by cell line.

Female euploid XX6 TBL cells however, secreted less hCG and PlGF, and fused at a lower rate than XX19/23 lines (Fig. S3B), pointing to differences among 46,XX euploid lines addressed below. Fusion indices were determined in unbiased fashion via computational analysis of DNA content (Hoechst staining), and revealed no other differences that rose to statistical significance. Likewise, differences in transwell migration rates missed the significance threshold (Fig. S3C), suggesting no overt karyotype-driven differences in this important trophoblast function (56, 57). All BAP-treated lines also gave rise to migratory cells that expressed the EVT marker HLA-G with similar, albeit variable frequency (Fig. S3D).

To develop a more comprehensive understanding for how monosomy X may impact TBL cell fates, we performed mRNA-seq in four independent rounds of BAP differentiation. We first assessed the BMP4-induced TF circuit triggered via *GATA2/3* and *TFAP2A/C* (20), all four of which were robustly expressed (>13 vst, variance-stabilizing counts, roughly approaching log2-scaled counts). XX19/23 and XY lines expressed moderately higher levels (log2FC 0.2-0.9, p.adj ≤ 0.05) of *TFAP2A, PGF* and *HLA-G* than their isogenic 45,X counterparts, which was not true for XX6 however (Fig. 2D). Of the TFs responding to the *GATA2/3 & TFAP2A/C* quartet (20), matching decreases in the male and female-derived panels were confined to a handful of transiently expressed TFs, except for *MEIS1* and *EPAS1*, which were modestly decreased in 45,X lines (Fig. S4A).

Next, we rigorously compared expression levels of lineage markers identified in single-cell RNA-seq studies of early human and macaque embryos (58–60). A subset of the human pre-gastrulation lineage markers (59) were re-classified recently based on single-cell RNA-seq from post-gastrulation macaque embryos (60) to resolve human TE, epiblast and amniotic lineages (61). As expected, we find that levels of both trophectoderm (TE)-associated gene sets (58, 61) significantly exceed levels of all other human lineage-associated gene sets in our BAP-treated cells (Fig. S4B,C) with a median differential of +1.5-2 vst counts (~4-fold difference, see methods). We also assessed all original gene sets (58–60) against the re-classified (TE, E-AM, and EPI) markers (61), and again find that both distinct TE gene sets far exceed levels of genes associated with all other lineages, and that early STB markers (59) represent the next-highly expressed set in our data (Fig. S4C). This is important in regards to a number of purported markers of the human amnion, which despite remaining poorly defined at present, have led to the suggestion that BAP treatment of primed hiPSC induces an amniotic rather than TE cell fate (62, 63). Yet, recent publications acknowledge that both naïve (+AP without BMP4) (62) and primed (36, 61) (+BAP) hiPSCs adopt TBL cell fates, and new work suggests that human trophoblast cells may differentiate through a transient amnion-like intermediate (61, 64). Our data across the full breadth of lineage-associated markers demonstrate that all of our BAP-treated lines reflect a predominantly TE and early STB-like expression profile (Fig. S4B,C).

Transcriptome-wide principal component (PC) analysis indicates that 45,X and euploid BAP-treated samples are clearly distinct, segregating along PC1 and PC2 largely by karyotype and donor, respectively (Fig. 2E). Surprisingly however, the XX6 euploid samples cluster with their otherwise isogenic 45,X samples (XO2/8/9), mirroring their relative decrease in hCG and PlGF levels (Fig. 2B,C). Because the predominant TS hypothesis posits a haplosinsufficiency of genes that escape XCI and were maintained on the Y, we next assessed allele-resolved and overall expression of PAR genes, interspersed X-Y gene pairs (“Pair”), and X-specific genes without Y-homolog. Overall, our phased variants covered 451 X-linked genes previously assessed in a large human GTEX study and meta-analysis (65), with sufficient allelic read depth to unambiguously call XCI status for up to 226 genes. A total of 166 genes were called (inactive/escape) across all 3 isogenic 46,XX lines, revealing excellent agreement overall (Fig. S5A). Of the 60 escapees we identified by allelic expression (lesser allele fraction, LAF ≥ 0.1, binomial p ≤ 0.05), 38 had previously been shown to escape XCI (65), and 4/22 remaining genes escaped across all three 46,XX lines (*POLA1, KLHL4, TMEM164, MBNL3*). Another 6/18 remaining genes escaped in at least two 46,XX lines (*FTX, SMS, AMMECR1, AMOT, STK26, IRAK1*), with the remainder reaching the escapee threshold in only a single line. Of the latter, only *MBTPS2* (in XX23), and *MID1, FAM199X, ELF4* and *MIR503HG* (in XX19) escaped partially (max. LAF ≤ 0.3) and were also overexpressed relative to 45,X samples. Three of these genes (*IRAK1, MBTPS2* and *SMS*) have been reported to variably escape in human placental samples (66, 67). Overall, these data indicate that aside from new escapee candidates and partial reactivation of at most four genes in one line, all three 46,XX lines faithfully maintained XCI over the course of these experiments.

These allele-resolved mRNA-seq data also reveal that the active X (Xa) in XX19 and XX23 is the same X retained in all female-derived 45,X lines, whereas this copy was chosen as the Xi in the XX6 line (Fig. S5B, see flipped A and B allele counts for each gene). Curiously, seven escapees common to XX19 and XX23 were expressed only from the Xa in the XX6 line, including validated escapees *MXRA5* (68), *PUDP* (69, 70), *STS* (71) and *SMS* (66), as well as *STK26, AMMECR1* and *AMOT* (72–74). We next queried levels of *XIST* and three other XCI-relevant non-coding RNAs to determine whether there was an association with this decreased level of escape in XX6 samples. As in hiPSC RT-qPCR (Fig, 1), *XIST* levels were higher in XX6 than XX19/23 TBL cells, but missed the significance threshold (Fig. 2F, p.adj = 0.22). There was no difference in two other X-linked non-coding RNA levels (*XACT, JPX*) amongst these female euploid TBL cells, but XX19/23-specific escapee *FTX* was expectedly higher than in XX6.

In standard differential expression, each of the XX19/23-specific escapees was also significantly lower in the XX6 line (p.adj ≤ 0.05, abs(log2FC) ≥ 0.3), whereas only XX6-specific escapee *SMC1A* was significantly higher (Fig. S5C). We also found many of PAR and X/Y pair (“Pair”) genes to be significantly decreased in XX6 relative to XX19 and XX23 euploid lines (Fig. 3A, S5C). Because the relative difference (log2FC panel and boxed vst differential heatmap) in these groups of escapees between XX6 and 45,X samples (XO2/8/9) was less pronounced than between XX19/23 and these X-monosomic lines (Fig. 3A, Fig. S5C), it was plausible that escapees across the XX6 Xi were repressed, rather than *cis*-acting variants reducing escape in each of these genes individually. In line with this interpretation, the X was significantly hypermethylated in XX6 hiPSC relative to XX19/23 lines (Fig. 3B), and >10% of X-linked promoter CpGs were differentially hypermethylated in the XX6 line (Fig. 3C,D). Importantly, this pattern most closely matches chromosomal probe density, and is significantly different (Kolmogorov Smirnov, KS test Fig. 3C) from the transition-specific trajectory we previously reported during Xi erosion (26). These data indicate that relative to XX19/C23, the XX6 Xi is hypermethylated (Fig. 3D, median +0.15 in DMP β value) across its entire length, including over differential escapees, and may suggest that variance in *XIST* levels contributes to variable escape. In summary, while differential and allelic expression analyses demonstrate that XCI remained virtually intact across all three female euploid TBL sets, XX6 samples revealed excessive repression of escapees across the Xi, relative to XX19 and XX23. Whether the mechanism of this repression rests on genetic variants boosting *XIST* expression, spreading or silencing will be addressed elsewhere.

**Figure 3:**
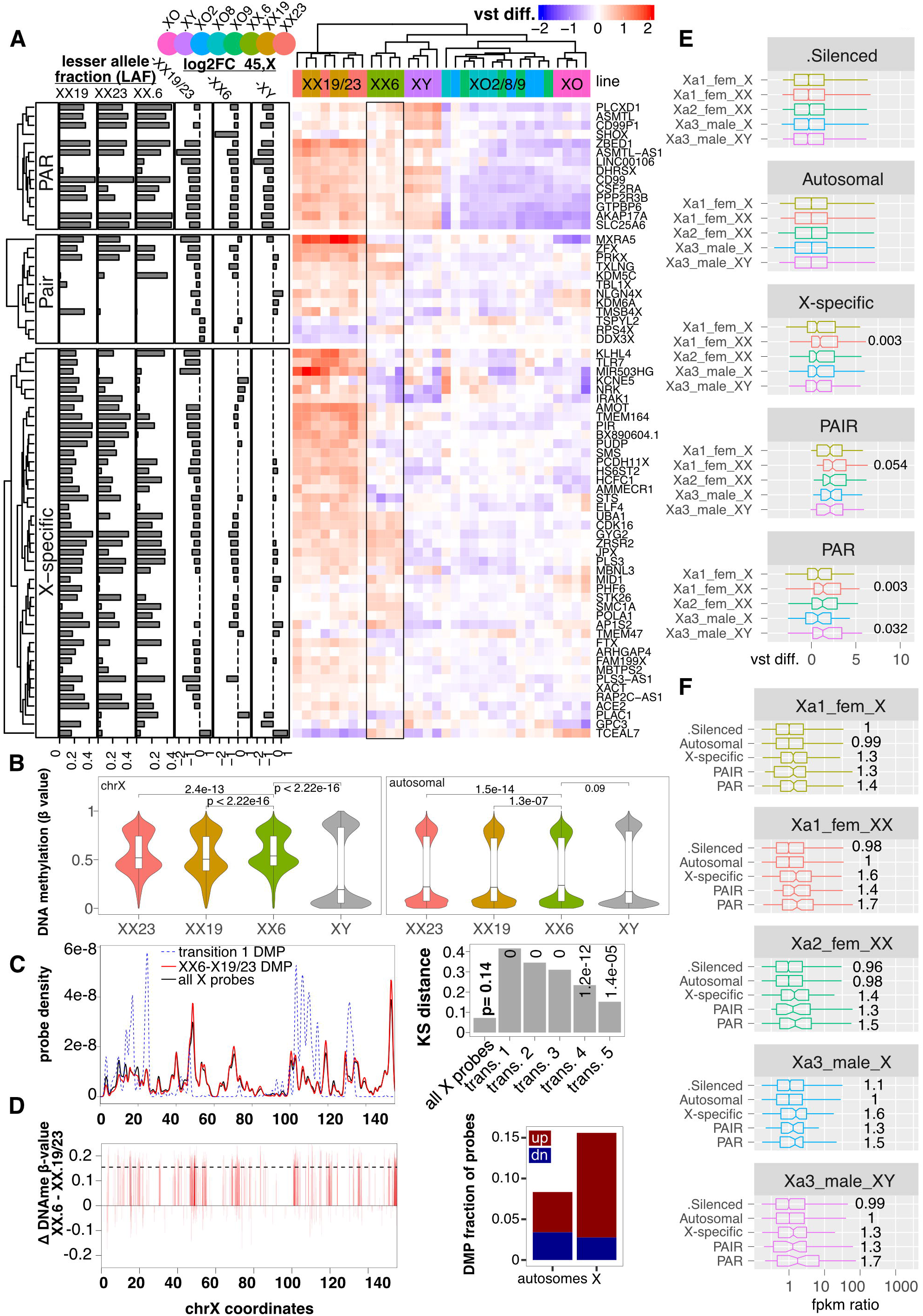
**A.)** Median-vst normalized expression values for PAR, Pair (X-linked X/Y pair gene) and X-specific escapee genes (as identified in Fig. S5). Heatmap columns denote lines, with XX6 expression values highlighted (black bordered box). Three left-most barplot panels report lesser allele fraction (LAF) as determined by phased RNA-seq. Three centered barplot panels report the estimated log2-scaled fold-change in expression (Log2FC) comparing 45,X samples to isogenic euploid controls (XX19/23 or XX6 for female-donor derived lines, or XY for male-derived lines). **B.)** Distribution of DNA methylation (β) values across X (top) and autosomes (bottom) in XX23, XX19 and XX6 lines, relative to published male control hiPSCs (118). Differences in median tested for significance by Mann-Whitney U test, with p-values listed above or below brackets. **C.)** *Left*: Chromosomal distribution of: differentially methylated probes (DMP) comparing XX6 to isogenic euploid XX19 and XX23 lines (red), DMPs previously identified in the first transition of Xi erosion (blue), and all probes on the MethylationEPIC array (black). *Right*: Kolmogorov–Smirnov (KS) test p-value and distance comparing XX6 DMP density across X to background distribution of all X-linked probes on the array, and Xi erosion transition-specific DMPs identified in (26). **D.)** *Left*: Difference in DMP β-value comparing XX19/23 to XX6 hiPSCs. Dashed line indicates the median change (β +0.15). *Right*: Fraction of DMPs with significantly greater (“up”) or lower (“dn”) β-value in XX6 relative to XX19/23 hiPSCs on X and autosomes. **E.)** Autosomal-median vst normalized expression of X-linked genes subject to XCI (‘Silenced’), relative to autosomal, X-specific escapees, PAR and X-linked PAIR genes, across all five conditions. Mann-Whitney U test p-value indicates significant difference from female-derived 45,X (“Xa1_fem_X”) samples. **F.)** Gene-length normalized expression (FPKM) comparing each class of genes in E.) to each other within each condition (see text). X:Autosome ratio for each class of genes denoted next to each boxplot.

The observation that XX6 TBL clustered with 45,X cells and phenocopied their significant decrease in hCG and PlGF secretion may reflect the consequences of their reduced escape from XCI, consistent with the haploinsufficient monosomy X hypothesis. We therefore included XX6 samples as a separate condition labeled by Xa identity (female-derived: Xa1 vs. Xa2, male: Xa3) throughout this analysis. Median expression of PAR, Pair and X-specific escapees in XX19 & XX23 (“Xa1_fem_XX”) but not XX6 (“Xa2_fem_XX”) is significantly higher than in female 45,X (“Xa1_fem_X”) lines, which is also true for PAR genes in the male panel (Fig. 3E, normalized to autosomal median). Comparing X:autosome (X:A) ratios across gene categories (Fig. 3F), we also find that genes subject to XCI are fully dosage compensated across male and female samples (X:A fpkm ratio of 1). This is consistent with the Xa hyperactivation hypothesis (75, 76), which posits that single-copy X-linked genes evolved to match transcription of autosomal genes that are expressed from two alleles. Interestingly, PAR, Pair and X-specific escapees are expressed at significantly higher levels (X:A fpkm ratio > 1), even when present in single-copy in 45,X samples. This result suggests that genes that evolved to escape XCI tend to also be expressed from the Xa well above average autosomal gene levels.

We then performed a systematic assessment of differentially expressed genes (DEGs), comparing: (1) isogenic X1 female euploid and 45,X samples (“fXO”), (2) isogenic male euploid and 45,X (“mXO”), and (3) a non-isogenic male euploid to female XO samples (“XY-fXO”). Altogether over 5000 genes were found to be differentially expressed (p.adj ≤ 0.05, abs(log2FC) ≥ 0.3) in at least two of these comparisons (Fig. 4A). As expected, these DEGs clustered samples by karyotype, but also featured highly significant overlap and concordance in direction (Fig. 4A), including 936 concordant DEGs out of 1283 common to all three comparisons (73% concordant, sign test p = 8.2×10^-285^). We next performed gene set enrichment analysis (GSEA), ranking genes by their individual fXO, mXO or XY-fXO DESeq2 Wald statistic, or the mean of their quantile-normalized scores (“aveXO”). As an additional control, we also ranked genes by the Wald statistic comparing karyotypically-identical 45,X samples across donors (“XOXO”). Expectedly, chrY and chrXp22 were recovered as significantly reduced in male XY-relative and all comparisons, respectively, alongside other chromosomal region-specific enrichments (Fig. S6). Among the computational modules of MsigDB, the placental gene module (#38) was the top gene set reduced in across all 45,X comparisons, and from a large human fetal single-cell RNA-seq dataset (77), three trophoblast-related gene sets are among the 15 most commonly reduced lineage-associated sets (Fig. S6). Interestingly, the Wikipathway (Fig. 4B), Reactome (Fig. S6), Hallmark and other MsigDB collections, point to impaired NRF2, cholesterol metabolism and estrogen signaling, all of which are important for placental function (71, 78–80). These terms were also significantly enriched in a recent transcriptome analysis of primary EVT and CTB (81), alongside gene sets relating to the cell cycle. Indeed, several of our significantly increasing terms related to the primary cilium (Fig. 4B: ‘Ciliopathies’, ‘Joubert Syndrome’; Fig. S6: ‘Anchoring of the Basal Body’, ‘BBSome-mediated cargo-targeting to cilium’ among others). The scale of this enrichment is clearly appreciable in the biological process (BP) and cellular component (CC) gene ontologies (GO), where proliferation, splicing and translation related categories are generally upregulated in 45,X TBL cells, but the two top terms (BP: ‘Cilium assembly’, ‘Cilium organization’, CC: ‘Centriole’ and ‘Ciliary basal body’) represent the primary cilium by some margin (Fig. 4C, Fig. S6). Among down-regulated gene sets, we find terms relating to the lysosome, autophagy, transmembrane transport, immunity, lipid and steroid metabolic processes among others (Fig. S6). The contrast between increased ciliary genes and decreased lysosomal genes in both female 45,X (Fig. 4D) and male 45,X (Fig. 4E) is particularly striking. Recent reports show the primary cilium is important for proper human trophoblast invasion and migration (82, 83). The primary cilium also has an inverse relationship with NRF2 (84–86), and NRF2 can activate PlGF expression directly (87). Indeed, NRF2 has been linked with birth outcomes in humans (78, 80) and mouse (79, 88–90). Likewise, cellular hallmarks of autophagy have been reported in late first trimester placenta (91, 92), and impaired autophagy has been implicated in recurrent and early miscarriage (93, 94).

**Figure 4:**
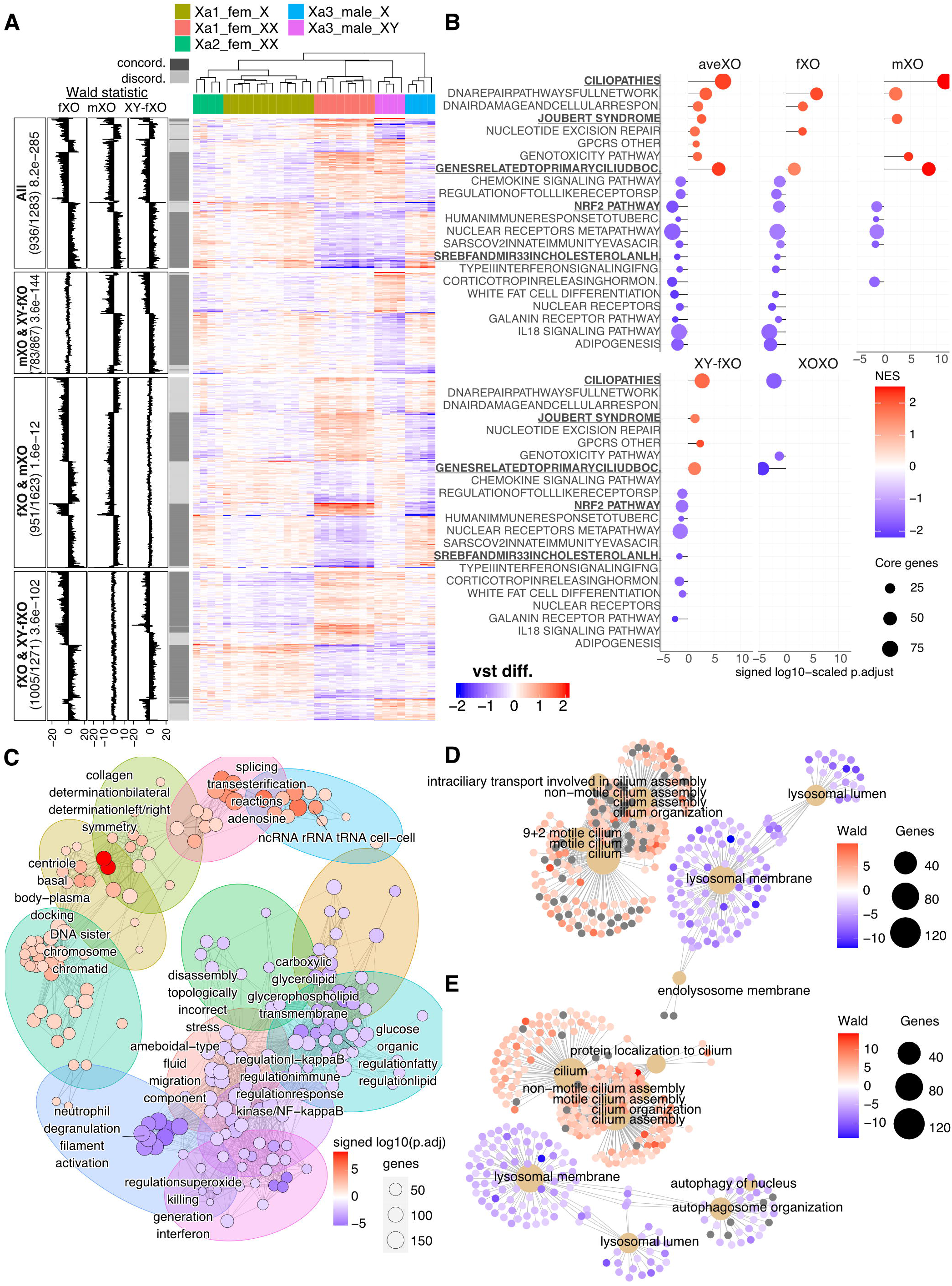
**A.)** Sample-level median-vst normalized expression (“vst diff.”) of all differentially expressed genes (DEGs, DESeq2 p.adj ≤ 0.05, abs(log2FC) ≥ 0.3) in 45,X lines relative to their isogenic euploid controls (“fXO”, mXO”), and between female-derived 45,X and male 46,XY replicates (“XY-fXO”). Number of overlapping and concordant DEGs identified in all three (top) or any two comparisons listed as a fraction alongside calculated p-value (sign test), left of barplot panels that report the DESeq2 Wald statistic. DEGs concordant across comparisons annotated in dark grey, discordant DEGs in light grey, in the heatmap-adjacent column. **B.)** Gene-set enrichment analysis (GSEA) against the Wikipathway collection (via MsigDB) for each comparison (“fXO”,”mXO”, “XY-fXO”), as well as a gene list re-ranked by the average Wald statistic from all three quantile-normalized sets (“aveXO”), and control comparison between male- and female-derived 45,X samples (“XOXO”). Bubble position, color and size, denote the signed log10-scaled GSEA p.adjust, the normalized enrichment score, and the number of core genes driving the enrichment, respectively, and are plotted opposite of abbreviated Wikipathway titles. **C.)** Semantic similarity-driven clusters and aggregated titles of biological process (BP) terms of the gene ontology (GO) enriched in GSEA (“aveXO”). Node colors and sizes denote signed log10-scaled GSEA p.adjust value and number of corresponding genes, respectively. **D.)** GSEA-enriched GO terms (beige) relevant to cilia, lysosomes, or autophagy and corresponding genes colored by “fXO” Wald statistic (red or blue for up- or downregulated DEGs in 45,X samples, grey for non-DEGs). **E.)** as in D.) for “mXO” Wald score-based GSEA terms.

To further clarify the relationships between these biological processes, we performed standard expression-trait correlation, and weighted co-expression network analyses (WGCNA) (95). First, we assessed how the levels of cell cycle and cell type markers (Fig. S4B,C), as well as PAR, Pair, X-specific, and all combined escapees (“All”), correlated with hCG and PlGF secretion (in matched RNA-seq & ELISA experiments, Fig. 2). “All” escapee, and PAR gene levels in particular, were highly and significantly correlated with hCG and PlGF secretion, as well as STB and EVT cell fates (Fig. 5A), whereas markers of cell cycle, CTB, and other proliferative cell fates correlated with each other. Unsurprisingly, STB and EVT cell fates anti-correlate with cell cycle and CTB markers, as STB and EVT arise from proliferative CTB but exit the cell cycle upon, respectively, fusion (96, 97) and maturation (81, 98).

**Figure 5:**
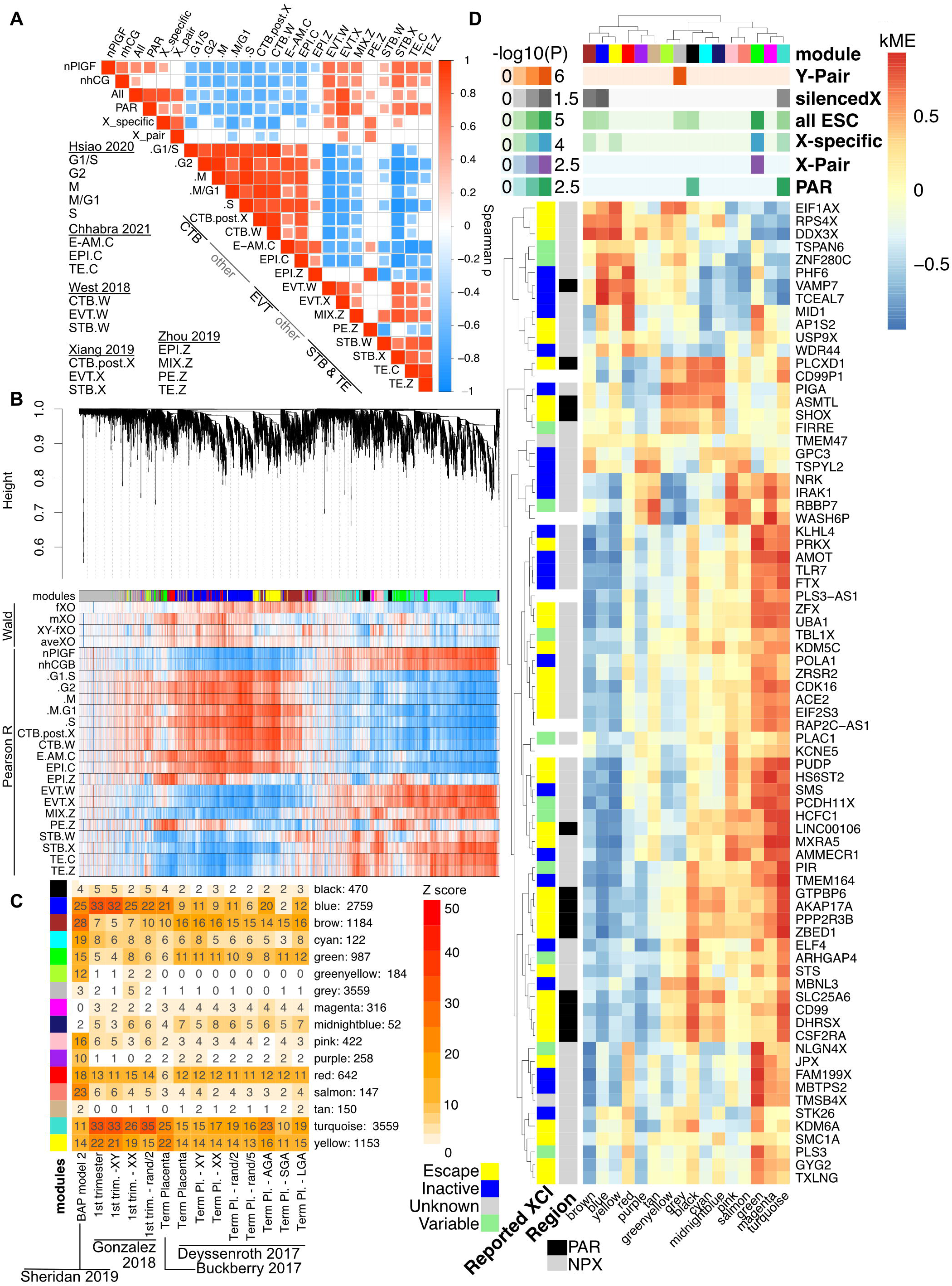
**A.)** Significant (p ≤ 0.05) expression correlation coefficients (ρ) between different escapee gene classes (All, PAR, X/Y-pair gene, and escapees lacking a Y homolog), and secreted PlGF and hCG levels, as well as cycling (119) and cell type markers from early human embryonic studies (58, 60, 61, 96), suffixed by first-author’s last initial (.W, .X, .Z, .C). Cell fates abbreviated for cyto-TB (CTB), early amnion (E-AM), epiblast (EPI), extravillous TB (EVT), mixed (MIX), primordial endoderm (PE) and syncytio-TB (STB). **B.)** Signed network from WGCNA plotted over module assignments, DESeq2 Wald stats (fXO, mXO, XY-fXO, and aveXO; up in red, down in blue), and Pearson correlation (r) with PlGF and hCG levels, as well as with median-normalized marker sets labeled as a in (A). **C.)** Permutation-based Z summary statistic for preservation of modules from B.) in another BAP dataset (99), (sex-stratified) first trimester chorionic villi sampling (CVS) (69), and two studies of term placenta (100, 101), the latter of which was further stratified by sex, birthweight or randomized (“rand”) to control datasets of similar size. **D.)** kME correlation coefficient of each escapee gene with its assigned module eigengene (heatmap), and enrichment analysis (log10-scaled p-value, Fisher test) for module assignment of various escapee gene classes. Region (PAR vs. NPX) and reported XCI status annotated on the left as in Fig. S5A.

Next, we performed WGCNA across all 32 BAP samples, which cleanly segregated by karyotype, except for XX6 samples that expectedly clustered with their 45,X counterparts (Fig. S7A). Plotted over DeSEQ2’s Wald scores, and gene-specific Pearson coefficients with hCG and PlGF levels (Fig. 5B), we observe a signed network of 16 modules that separates genes into two groups: Group 1 genes (left side) largely correlate with DEGs increasing in 45,X samples and anti-correlate with hCG and PlGF secretion, whereas group 2 genes (right side) co-correlate with DEGs decreasing in 45,X and hCG and PlGF levels. Strikingly, group 1 genes correlate with cell cycle and CTB markers and anti-correlate with STB and EVT markers, whereas the inverse is true for group 2 genes. These data suggest that the network is shaped by mutually exclusive cell fates. To determine which specific modules were significantly driven by the contrast between euploid vs. 45,X expression, we quantified the degree and significance of their preservation in a subset of exclusively 45,X samples and a mixed control dataset of equal size. Preservation (Z) scores of modules representing over two-thirds of all genes show a decrease by 20-80 standard deviations in the 45,X-only dataset relative to the mixed karyotype set (Fig. S7B). Correlating each module to traits of interest (Fig. 5A), we find the same set of modules to be strongly anti- or co-correlated with euploidy, PAR expression, hCG & PlGF secretion (Fig. S7C).

To test whether these modules are recovered in independent BAP and primary placental samples, we performed module preservation analysis against RNA-seq datasets from another hiPSC-based BAP model, for pre-eclampsia (99), first trimester chorionic villi samples (CVS) (69), and two WGCNA studies on placental samples at term (100, 101). Remarkably, the same 45,X – euploid contrasting modules (Fig. S7B,C) are also moderately to highly preserved in primary placental samples, irrespective of (fetal) sex or birth weight categories (Fig. 5C). Because higher CTB and cycling markers correlated with monosomy X in our WGCNA modules (Fig. S7C) and standard correlation analysis (Fig. 5A), we interpret this high level of preservation to reflect variable cell type composition (eg. CTB vs. STB) that is inherent in the sampling of first trimester CVS, and term placenta. Sampling of any primary tissue is necessarily variable in cell type composition, and this variance is frequently captured in WGCNA (102). Here, in the context of our BAP model, gene modules preserved in CVS and placental samples may suggest that monosomy X hinders or delays commitment of cycling BAP-derived CTB-like cells to post-mitotic STB and EVT cell fates, thereby increasing CTB marker representation and continued expression of cycling markers. Indeed, our WGCNA modules are most strongly preserved in the first-term trimester CVS, in which proliferation and cell fate commitment are likely even more variable than in term placenta (Fig. 5).

We also tested whether modules were over-represented for gene sets assessed in the differential expression analysis. We recovered many similar terms (Fig. S8) relating to cell cycle and primary cilium (blue, yellow), translation, autophagy and metabolism (brown, yellow), membrane-anchored signaling pathways (green), immune regulation (midnightblue), adipogenesis and the lysosome (turquoise), Among the human fetal single-cell cell type terms, we again find three trophoblast gene sets (brown, turquoise), overall indicating strong overlap with enriched terms from differential expression (Fig. 4, S6), but providing module-level resolution of cellular functions.

Finally, to implicate the dosage of specific X/Y-linked gene classes in TBL differentiation, we first tested which modules featured an over-representation of PAR, Pair or X-specific escapees. The green module was significantly enriched (p ≤ 0.05) for all escapees as one class (“allESC”), escapees without Y-homolog (X-specific), and X-linked X/Y pair genes, whereas PAR genes were most over-represented in the black and turquoise modules. This latter module was of particular interest because it was also highly enriched for genes from MsigDB’s placental gene module (#38) and human fetal EVT markers (Fig. S8), correlated strongly with TE, STB and EVT markers identified across numerous early human embryonic studies, as well as hCG and PlGF levels, and best reflected euploidy and escape from XCI (Fig. 5B, S7C). Importantly, the turquoise module was also the most preserved in first trimester and term placental RNA-seq samples (Fig. 5C).

To identify potential X-linked drivers strongly associated with specific modules, we determined the degree of correlation between each individual gene with its module eigengene (averaged module expression profile across samples). Raising this coefficient (kME) to the same power as the network, and plotted over the degree of connectivity between genes, we find that PAR gene *ZBED1* is the top X/Y-linked hub gene in the turquoise module, ranking 272th of 3559 genes (kME = 0.92) in the module overall. Additionally, other PAR genes (*PPP2R3B, GTPBP6, AKAP17A* and *CD99* > 0.8 kME) and escapees repressed in XX6 ranked highly in this module (kME 0.86 – 0.7 for *AMOT, PUDP, SMS & STS*). In summary, our WGCNA analysis indicates that PAR expression most strongly reflects placental gene expression in the TBL (BAP) model, and may serve to prioritize a core set of X/Y-linked genes for follow-up in other *in vitro* models of human extra-embryonic development.

## DISCUSSION

As a leading cause of spontaneous termination in humans (15), monosomy X can serve as a penetrant genetic model for miscarriage. Few genome-wide association studies on spontaneous or recurrent miscarriage have been published to-date (103–105), which face the additional challenge of accounting for and excluding embryonic/fetal karyotypic changes (106, 107). Characterizing the impact of monosomy X using *in vitro* human cell models may therefore provide a complementary approach towards implicating cellular functions and pathways in miscarriage.

The hiPSC BAP model (18) used herein has previously revealed sex-divergent expression patterns (46), addressing another important question in trophoblast biology (108–110). In contrast, we applied this model to identify TBL cellular phenotypes and expression signatures that are common to the absence of the Xi or Y (rather than sex-divergent), to better understand the consequences of this compound haploinsufficiency. We also rigorously validated the BAP model by comparing expression of markers identified across a range of independent early human and primate embryonic studies (58–61, 96), to find TE and STB markers to be the predominantly expressed lineage-associated gene sets in our experiments (Fig. S4).

Importantly, we find secretion of STB-produced hCG and PlGF is significantly decreased in 45,X cells compared to isogenic euploid controls (Fig. 2B,C). Although differences in STB fusion index or the fraction of HLA-G^+^ cells did not rise to statistical significance (Fig. S3), *PGF* and *HLA-G* transcript levels, respective markers of STB and EVT, were also reduced significantly in 45,X samples (Fig. 2D). This is relevant because misregulation of HLA-G alone can result in miscarriage (111), and significantly lower PlGF levels have previously been reported in X-monosomic first trimester pregnancies (112, 113).

Two related insights emerged from the larger pattern of global 45,X-associated expression changes: 1.) Among significantly enriched gene sets undergoing concordant changes in male -and female-derived 45,X samples (Fig. 4), we find increased proliferation-associated terms (primary cilium, DNA replication, splicing, translation) and decreased terms related to maturing STB and EVT cellular functions (transmembrane transport, immune-regulation, and metabolism), which included lysosomal processes like autophagy. 2.) Likewise, standard correlation and WGCNA reveals cell cycle and CTB markers correlate with genes that increase in 45,X samples, whereas STB and EVT markers correlate and cluster with genes that decrease in 45,X relative to euploid TBL cells (Fig 5). Because both STB and EVT cells derive from CTB, but must exit the cell cycle upon fusion or maturation, the correlation between monosomy X and cell cycle / CTB markers (Fig. 5A,B, S7D) suggests 45,X TBL cells are still skewing towards actively cycling CTB at the end of the 8-day BAP differentiation, which may explain their lower secretion of hCG and PlGF (Fig. 2).

While the molecular basis of this delay in committing to STB or EVT cell fates remains unclear, our WGCNA indicates that PAR genes in general, and *ZBED1* specifically, are strongly positively correlated and well connected inside the turquoise placental gene module (Fig. 5D, S7C,D). This module was also the top preserved gene module in RNA-seq studies of term placenta and especially first trimester CVS (Fig. 5C), which are likewise heterogenous in respective CTB vs. STB and EVT contributions due to sampling. Intriguingly, *ZBED1* does regulate proliferation (114) and is expressed in human placenta, with higher levels in post-mitotic STB expression than CTB (115).

Our study highlights promising areas for follow-up in future *in vitro* work and study of primary samples. For example, it is unclear whether higher expression of primary cilia components merely reflects the higher frequency of ciliary re-synthesis in cycling 45,X cells, or altered function of this important signaling organelle. While mouse trophoblasts lack cilia, human trophoblast carry cilia (82, 83), and cilia regulate autophagy and NRF2 (86, 116). Curiously, lysosomal genes were widely down-regulated in 45,X BAP samples (Fig. 4), and impaired autophagy has previously been implicated in recurrent miscarriage (92–94, 117). Whether placental autophagy is de-regulate in miscarried 45,X conceptuses specifically, and whether cell fate proportions are altered in such 45,X-associated CVS or placental samples, would therefore be of particular interest towards understanding why human monosomy X terminates early.

## MATERIALS AND METHODS

### Reprogramming and iPSC culture

Fibroblasts were reprogrammed into hiPSCs at the UConn Stem Cell Core, using the CytoTune iPSC 2.0 Sendai Reprogramming kit (Thermo Fisher Scientific, Waltham, MA). All hiPSC clones were initially cultured on mitotically inactive mouse embryonic fibroblasts (MEFs) in standard human iPSC media (80% DMEM/F12, 20% Knockout Serum Replacement, 1% Glutamax, 1% Non-Essential Amino Acids, 0.1% β-Mercaptoethanol, and 8ng/mL FGF), and maintained by weekly mechanical passaging. Subsequently, hiPSCs were transitioned to mTESR media (Stem Cell Technologies, Cambridge, MA), grown on extracellular matrix (Geltrex, Thermo Fisher, Waltham, MA), and passaged weekly with EDTA.

### Cytogenetic analysis and DNA methylation profiling

Karyotype analysis was performed on 20 GTG banded metaphase cells at the UConn Center for Genome Innovation. Cytogenomic analysis was performed at the UConn Center for Genome Innovation on the CytoSNP-850k v1.2 (Illumina) platform, using phenol-chloroform extracted genomic DNA.

For DNA methylation analysis, genomic DNA from euploid 46,XX cells was bisulfite-converted using the EZ DNA Methylation Kit (Zymo Research, Irvine, CA), labeled and hybridized using the Infinium Methylation EPIC BeadChip Kit (Illumina, San Diego, CA) following standard protocol of each manufacturer, and scanned on a NextSeq 550 system. The data were analyzed using the minfi R package with IlluminaNormalization (120). Probes with a UCSC_RefGene_Group designation of TSS1500, TSS200, or 5_UTR were designated as promoter probes. Differentially methylated probes (DMPs) characterized previously (26) and associated with specific transitions during Xi erosion were queried (in Fig. 1F). New DMPs distinguishing XX6 from XX19/23 were called via minfi (p-value ≤ 0.05). KS distances and KS test significance were calculated comparing the gaussian probe densities across the X chromosome for each probe set (all X-linked Infinium MethylEPIC probes, and transitions 1-5 from (26)). Change in DNA methylation (β value) was calculated as the difference between the mean XX6 probe β and the mean XX19/23 probe β value.

### DNA sequencing and phasing

High molecular weight (HMW) genomic DNA was prepared following cell lysis in 10% sarcosyl/5uM NaCl/100uM EDTA/100uM Tris pH 8 with 1mg/mL Proteinase-K, and incubation at 55°C overnight. After RNA digestion (20ug/mL RNase-A) for 30 minutes at 37°C, HMW genomic DNA was isolated by phenol chloroform extraction in Phase-Lock Gel Heavy tubes (Quantabio, Beverly, MA), and precipitated in 70% ethanol, washed in 70% ethanol, and re-solubilized in 10 mM Tris pH8, 0.1 mM EDTA (Te). The HMW-gDNA was sequenced to ~30x coverage, following library preparation on the 10X Genomics Linked-Read platform at the UConn Center for Genome Innovation. LongRanger (10X Genomics) was used for read alignment and phasing of variants genome-wide. X-linked phased variants were supplied alongside RNA-seq data to obtain A and B allele counts for X chromosome genes using phASER (121).

### RT-qPCR

Reverse transcription was performed using the iScript gDNA Clear cDNA Synthesis Kit (Bio-Rad, Hercules, CA). Quantitative PCR was performed on the resulting cDNA using the iTaq Universal SYBR Green Supermix (Bio-Rad, Hercules, CA) with the primers XIST_F: CTCCAGATAGCTGGCAACC; XIST_R: AGCTCCTCGGACAGCTGTAA; GAPDH_F: CTGGGGCTGGCATTGCCCTC; GAPDH_R: GGCAGGGACTCCCCAGCAGT.

### Trophoblast Differentiation

TBLs were differentiated from hiPSC as described (18) with minor modifications. Briefly, confluent hiPSC clones cultured in mTeSR were dissociated with Accutase and plated at 50,000 cells/well of a 6well plate in mTeSR with 10uM Y-27632 (Tocris Bioscience, Bristol, UK) for one day. Then media was changed to mouse embryonic fibroblast conditioned media (MEF-CM), supplemented with 8ng/uL human basic FGF (Thermo Fisher Scientific) and 10uM Y-27632. The following day, media was switched to BAP differentiation media, which consisted of MEF-CM with 10ng/mL BMP4 (Peprotech, Rocky Hill, NJ), 1uM A83-01 (Stem Cell Technologies) and 0.1uM PD173074 (Stem Cell Technologies). Media was changed daily until day 8 when cells and supernatant were harvested for RNA collection, IF, or ELISA, respectively. For ELISA, supernatants were diluted 1:1,000 for the hCG ELISA (GenWay Biotech, San Diego, CA) and 1:50 for the PIGF ELISA (R&D Systems, Minneapolis, MN).

### Immunocytochemistry

Immunocytochemistry was performed on the hiPSCs using the PSC-4 Marker Immunocytochemistry Kit (Thermo Fisher Scientific, A24881). For immunocytochemistry of trophoblast markers after eight days of BAP differentiation, cells were fixed in 4% PFA for 30 minutes at 4°C, washed with 0.1% Triton-X, permeabilized with 0.5% Triton-X for 5 minutes. Following blocking in 5% normal goat serum/2% BSA/0.1% Tween-20, cells were incubated overnight at 4°C with rabbit monoclonal ATPase antibody (Abcam, ab76020, 1:500), mouse monoclonal HLA-G antibody (Abcam, ab52455, 1:500) or mouse hCGb antibody (ThermoFisher, MA-35020, 1:100). Slides were then washed twice in 0.1% Tween-20, incubated with AlexaFluor-555 Goat-anti-Mouse and AlexaFluor-647 Goat-anti-Rabbit Secondary Antibodies (Thermo Fisher Scientific, Waltham, MA, both at 1:500) for 1 hour at room temperature. Cells were washed twice in 0.1% Tween-20, stained with Hoecsht-33342, and mounted with ProLong Gold Mounting Medium. Fluorescent images were taken on an EVOS Auto FL Cell Imaging System (Thermo Fisher Scientific, Waltham, MA).

### Fusion Index

The fusion index was calculated by automated analysis of Hoechst-stained images using CellProfiler (122). Nuclei were identified as objects, and recorded by their integrated intensity and size, using hiPSCs to provide an empirical null distribution representing unfused cells. Using the 99^th^ percentile of the null distribution as an integrated intensity cutoff, TBL nuclei with greater or equal to 99^th^ percentile intensity were designated as fused. This threshold was additionally normalized by the difference in median intensity to account for differences in staining or illumination. The fusion index was calculated by dividing the integrated intensities of all fused objects by the total integrated intensity.

### Transwell Migration Assay

The trophoblast differentiation was performed in Corning BioCoat Matrigel Invasion Chambers (Corning, NY) with 8 μm pores. 50,000 iPSCs were plated on the top of each chamber and differentiated, fixed and stained as described above. The migration index was calculated by dividing the total DNA integrated intensity of the bottom by the sum of integrated intensity on the top and bottom of the transwell.

### RNA Sequencing

RNA was extracted from iPSCs using the PureLink RNA Mini Kit (Thermo Fisher Scientific, Waltham, MA). For 3’mRNA-seq, libraries were prepared using the Quant-seq 3’ mRNA Library Prep Kit FWD (Lexogen, Greenland, NH) and single-end 75bp reads were sequenced on the NextSeq 500 (Illumina, San Diego, CA). Genes queried for hiPSC identity were listed on the TaqMan hPSC Scorecard Assay (Thermo Fisher Scientific, Waltham, MA). For standard mRNA-seq, libraries were prepared at the UConn Center for Genome Innovation using the Illumina Strand mRNA Kit and 100bp paired-ends reads were sequenced to an average depth pf 40 million reads/replicate on the NovaSeq (Illumina, San Diego, CA).

### Allelic RNA-seq, differential expression, GSEA and WGCNA

Read pairs were trimmed using cutadapt v2.7 (123), aligned to the human genome (hg38) with hisat v2.2.1 (124), and quantified against GENCODE version 36 (125), with featureCounts v2.0 from the Rsubread package (126). For escapee calls based on phased linked read variants from linked-read sequencing, A and B allele counts from phaser (121) were tabulated by cell line across replicates, and 46,XX allelic counts were adjusted for each gene based on the absent allele count in 45,X replicates and relative total allelic read depth in 45,X and 46,XX replicates. Escapees calls were made using a binomial test (lesser allele fraction, LAF > 0.1, p ≤ 0.05) for all X-linked genes with a minimal allelic read count to identify escapees with LAF ≥ 0.2 (power of 0.9, given a read error rate of 0.01).

For differential expression using DESeq2 (127), count tables were filtered for genes with sufficient expression (10 count average, with 20 counts in at least 2 samples). Surrogate variables were estimated using the sva package (128), and added to the DESeq2 design, which compared conditions based on Xa identity (female-derived: Xa1, Xa2, male-derived: Xa3) and monosomy X (45,X vs. 46,XX or 46,XY). Gene-set enrichment analysis (GSEA) using clusterProfiler (129) was performed on all genes ranked by DESeq2’s Wald statistic in three separate conditions, as well as the average of their quantile-normalized Wald scores to ensure equal weighting. GSEA terms were abbreviated and plotted using clusterProfiler, across a range of p.adj thresholds (all ≤ 0.1 in the aveXO GSEA) to accommodate plot size limitations. Weighted gene co-expression network analysis (WGCNA) was performed on vst counts for all genes with Entrez geneid (130), using the WGCNA package (95), as a signed hybrid network using the biweight midcorrelation raised to a soft thresholding power of 16 (scale-free topology fit ≥ 0.85). Modules were correlated to normalized hCG and PlGF ELISA values, and to averaged early embryonic lineage marker sets, which were median vst normalized to ensure equal weights across all sets. Module preservation analysis was performed with the WGCNA package against published BAP (99), CVS (69) and term placenta (100, 101) RNA-seq datasets. Enrichment analysis of escapee gene class across WGCNA modules applied a hypergeometric test (p ≤ 0.05). Enrichment analysis of gene sets across WGCNA modules was performed using the compareCluster function of the clusterProfiler package (129).

## ACKNOWLEDGMENTS

We would like acknowledge Yaling Liu and Leann Crandall at UConn Health’s Cell and Genome Engineering Core for hiPSC reprogramming and immunocytochemistry services. We would also like to thank Lisa LaBelle and Judy Brown at the UConn Chromosome Core for cytogenetic characterization of our hiPSC lines services, and Bo Reese at the UConn Center for Genome Innovation for mRNA-seq library preparation and sequencing. This work was supported by NIH grant R35GM124926 to S.F.P.

## Notes

### Competing Interest Statement

The authors have declared no competing interest.

