## Supplementary Figures for "Monosomy X in isogenic human iPSC-derived trophoblast model impacts expression modules preserved in human placenta"

**SUPPLEMENTARY MATERIAL**

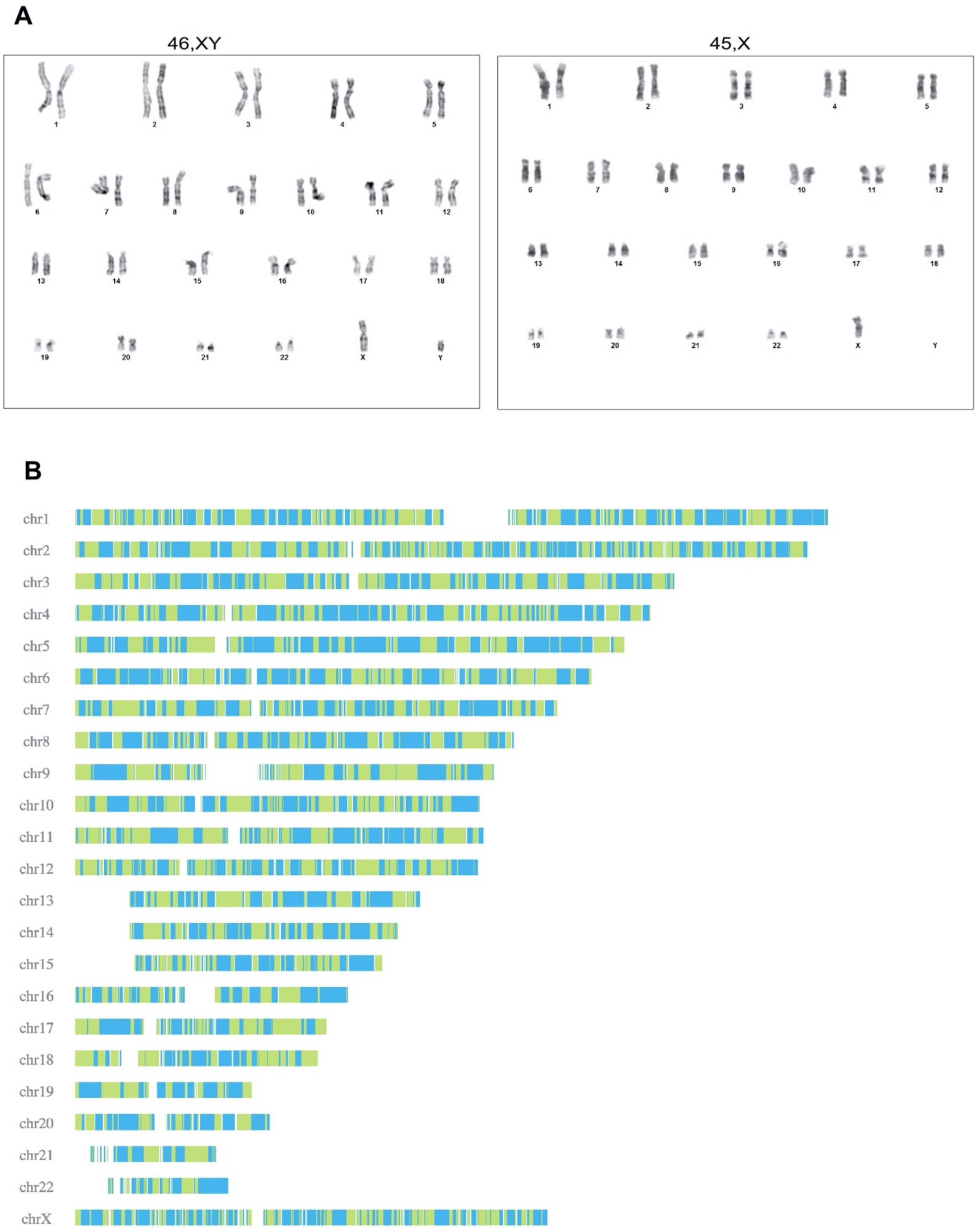


**Figure S1**

**Figure S1: A.)** Karyotype images of isogenic 46,XY (left) and 45,X (right) hiPSC clones from the male donor. **B.)** Phaseblock map from linked-read sequencing (10X Genomics) of euploid 46,XX hiPSCs from the female donor.

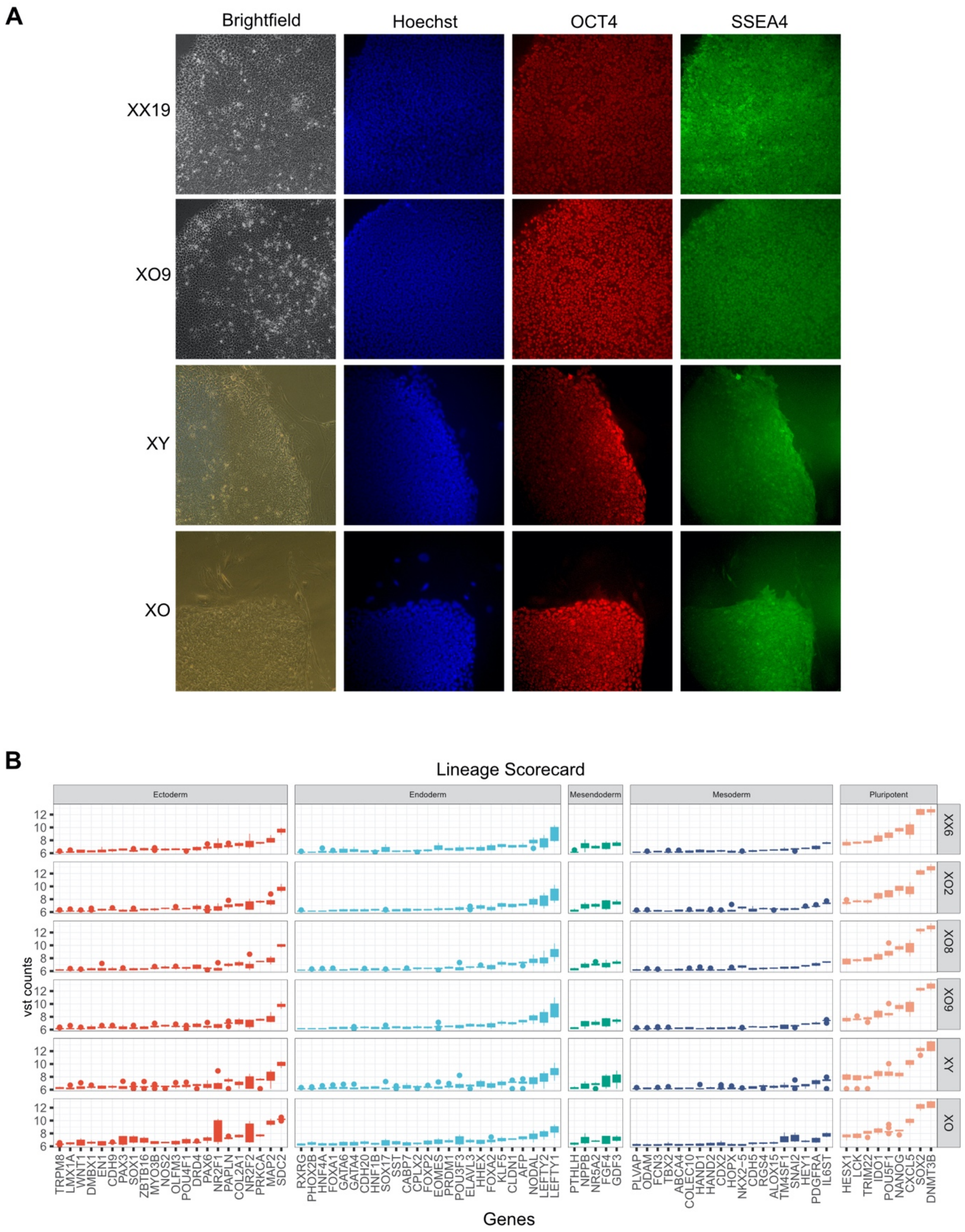


### Figure S2

**Figure S2: A.)** Representative brightfield and immunocytochemistry images of euploid and 45,X iPSC clones. **B.)** Variance-stabilized-transformed (vst) counts from 3' mRNA sequencing of ectoderm, endoderm, mesendoderm, mesoderm, and pluripotency genes in hiPSCs lines.

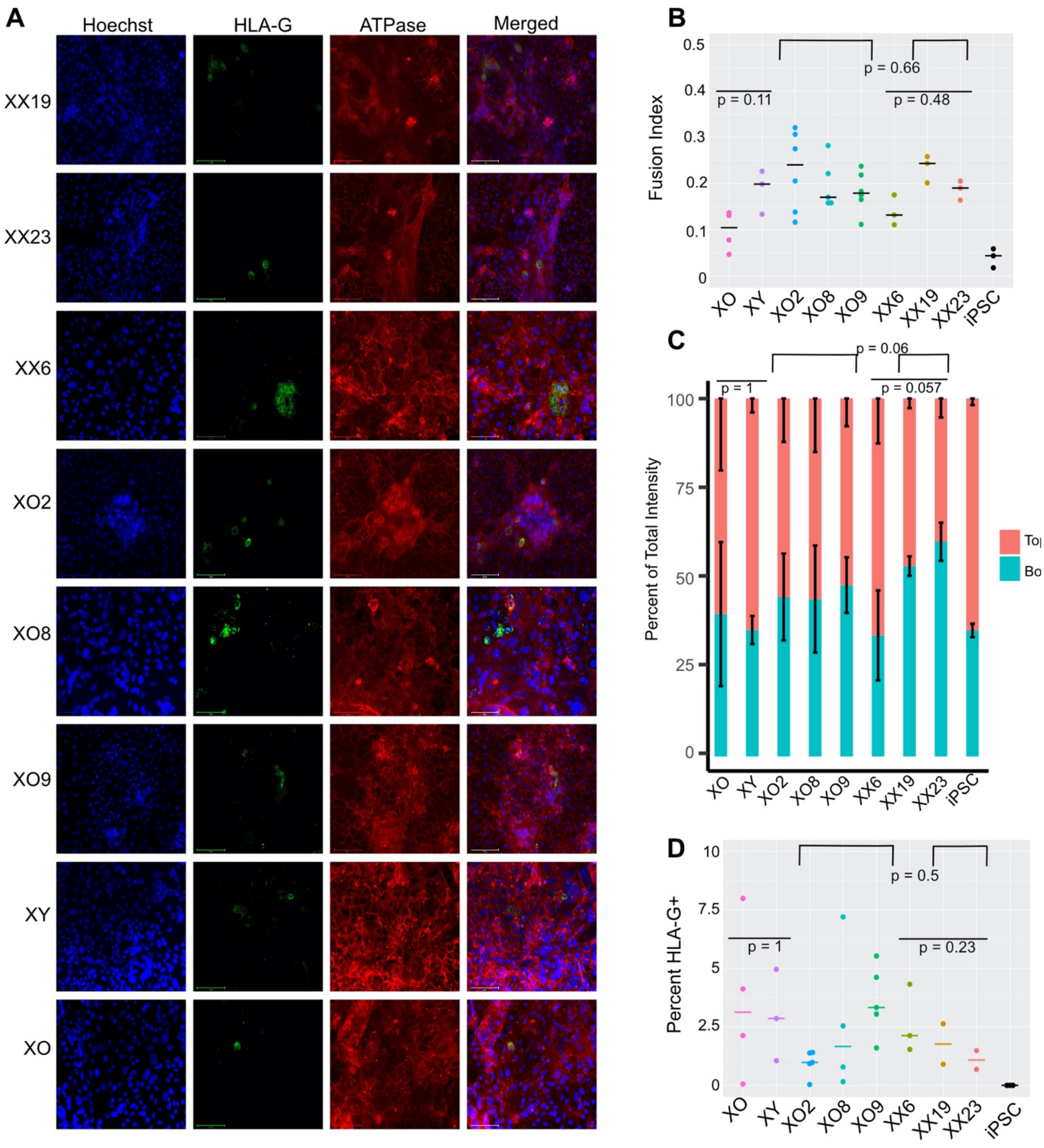


**Figure S3**

**Figure S3: A.)** IF images of TBLs stained for HLA-G and Na+/K+ ATPase, nuclei counterstained with Hoechst. **B.)** Fusion index in TBLs, calculated as the ratio of contiguous nuclear DNA signal over total DNA. Mann-Whitney-U p-values compare 45,X to otherwise isogenic euploid controls (as denoted by brackets). **C.)** Transwell top and bottom -localized DNA signal as a function of total DNA intensity across both sides. **D.)** Percentage of HLA-G+ cells with P-values from Mann-Whitney-U tests comparing 45,X to otherwise isogenic euploid controls (as denoted by brackets).

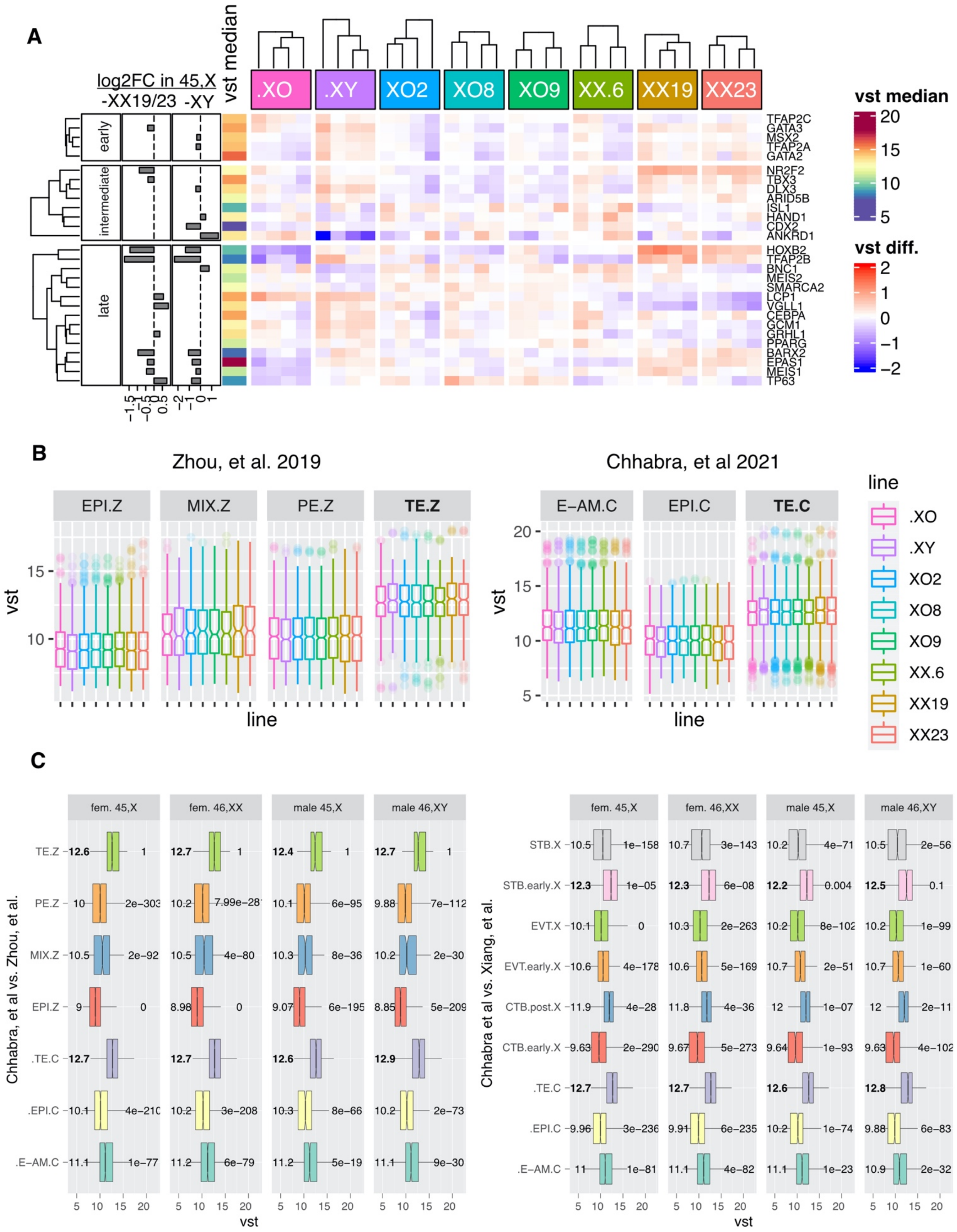
A
log2FC in 45,X
-XX19/23 -XY
vst median
.XO
.XY
XO2
XO8
XO9
XX.6
XX19
XX23
early
intermediate
late
TFAP2C
GATA3
MSX2
TFAP2A
GATA2
NR2F2
TBX3
DLX3
ARID5B
ISL1
HAND1
CDX2
ANKRD1
HOXB2
TFAP2B
BNC1
MEIS2
SMARCA2
LCP1
VGLL1
CEBPA
GCM1
GRHL1
PPARG
BARX2
EPAS1
MEIS1
TP63
vst median
vst diff.
B
Zhou, et al. 2019
Chhabra, et al 2021
EPI.Z
MIX.Z
PE.Z
TE.Z
E−AM.C
EPI.C
TE.C
vst
line
C
fem. 45,X
fem. 46,XX
male 45,X
male 46,XY
Chhabra, et al vs. Zhou, et al.
Chhabra et al vs. Xiang, et al.
TE.Z
PE.Z
MIX.Z
EPI.Z
.TE.C
.EPI.C
.E−AM.C
STB.X
STB.early.X
EVT.X
EVT.early.X
CTB.post.X
CTB.early.X
vst

**Figure S4**

**Figure S4: A.)** Absolute expression (vst) of BMP4-induced transcription factors (TFs) reported prior (1), across all BAP-treated replicates for each line. **B.)** BAP sample expression (vst) levels for markers of epiblast (EPI.Z), mixed lineage (MIX.Z), primordial endoderm (PE.Z) and trophectoderm markers (TE.Z) reported in (2), as well as a re-classified (3) subset markers for the early amnion (E-AM.C), EPI.C and TE.C. **C.)** *Left:* Distribution of BAP-treated expression (vst) levels for unified but distinct sets of markers from two studies in B.). *Right:* as on the left but for a unified and distinct set of re-classified (3) and original (4) early human embryonic lineage markers (suffixed '.X'). Median vst denoted on the left of each boxplot, Mann-Whitney U test p-value relative to TE.C marker set on the right of each boxplot. Each set of markers was independently assessed across all four karyotypes (panel titles).

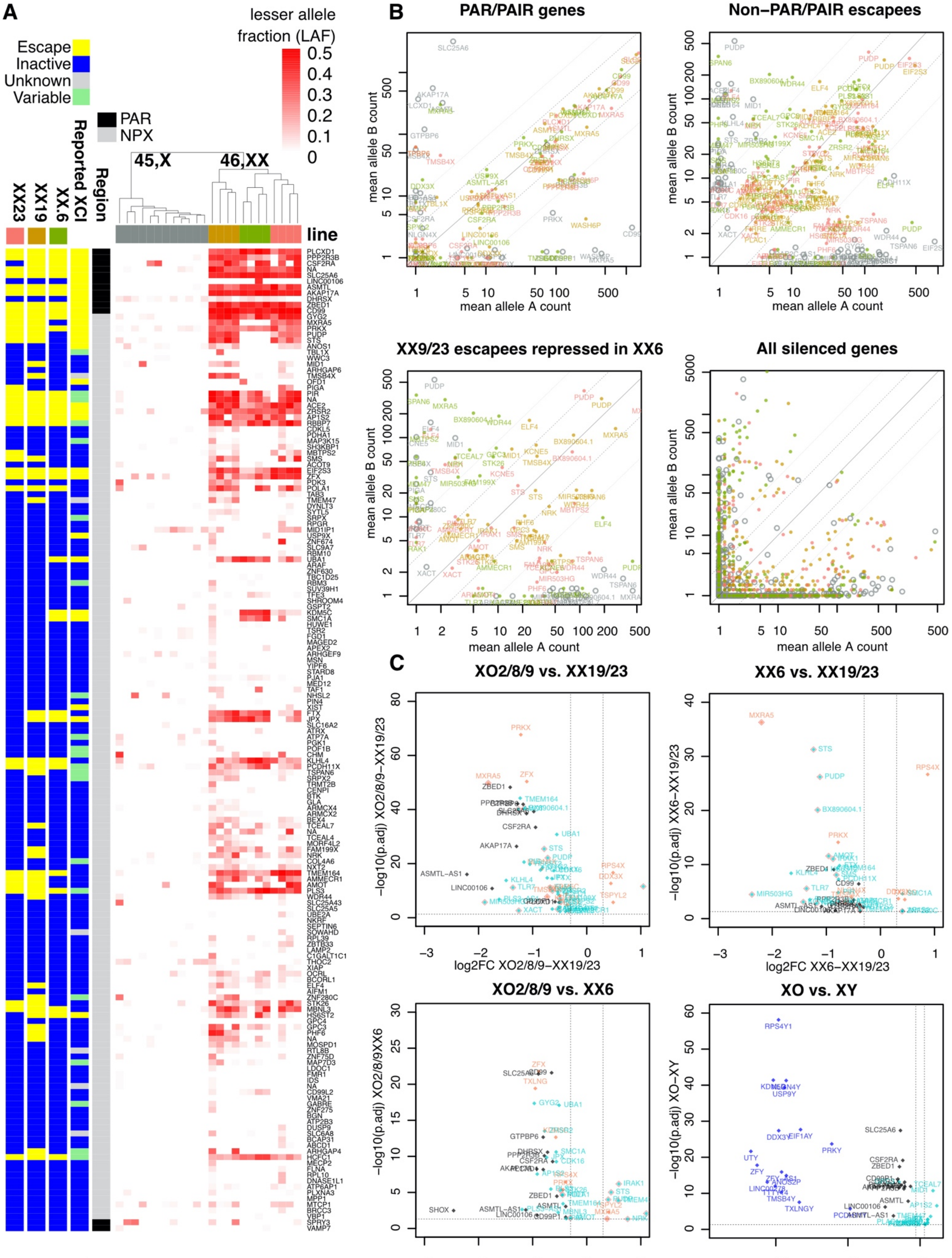
A
Escape
Inactive
Unknown
Variable
lesser allele fraction (LAF)
PAR
NPX
45,X
46,XX
XX23
XX19
XX.6
Reported XCI
Region
line
B
PAR/PAIR genes
Non-PAR/PAIR escapees
XX9/23 escapees repressed in XX6
All silenced genes
mean allele A count
mean allele B count
C
XO2/8/9 vs. XX19/23
XX6 vs. XX19/23
XO2/8/9 vs. XX6
XO vs. XY
log2FC XO2/8/9−XX19/23
log2FC XX6−XX19/23
log2FC XO2/8/9−XX6
log2FC XO−XY

**Figure S5**

**Figure S5: A.)** Lesser allele fractions (LAF) for X-linked genes from allele-specific RNA-seq data in 45,X and 46,XX TBL samples. Reported XCI status from (5), chromosomal region (PAR/NPX), and independent calls for genes escaping (yellow) or subject to XCI (blue) indicated on the left. **B.)** Gene-level allelic RNA-seq counts phased to A & B alleles on log-scaled plot, with 46,XX lines colored as in A.). Diagonal lines indicate level of escape relative to Xa expression (center line = 50% of allelic counts from Xi, others 33% and 10% from the Xi). Panels split between PAR/PAIR genes, escapees without Y homolog (non-PAR/PAIR), escapees repressed in XX6, and all genes subject to XCI. **C.)** Differential expression volcano plots of log10-scaled p.adj (DESeq2) over log2FC estimate for PAR (black), PAIR (orange) and other escapees without Y homolog (turquoise), bordered in red for XX19/23 escapees repressed in XX6 samples. Panels split DEGs in 45,X lines relative to either isogenic XX19/23 or XX6 euploid samples, compare XX6 to XX19/23, or male-derived 45,X and 46,XY lines (Y homologs in blue). Dashed lines denote differential expression thresholds ($p.adj \leq 0.05$, $abs(log2FC) \geq 0.3$).

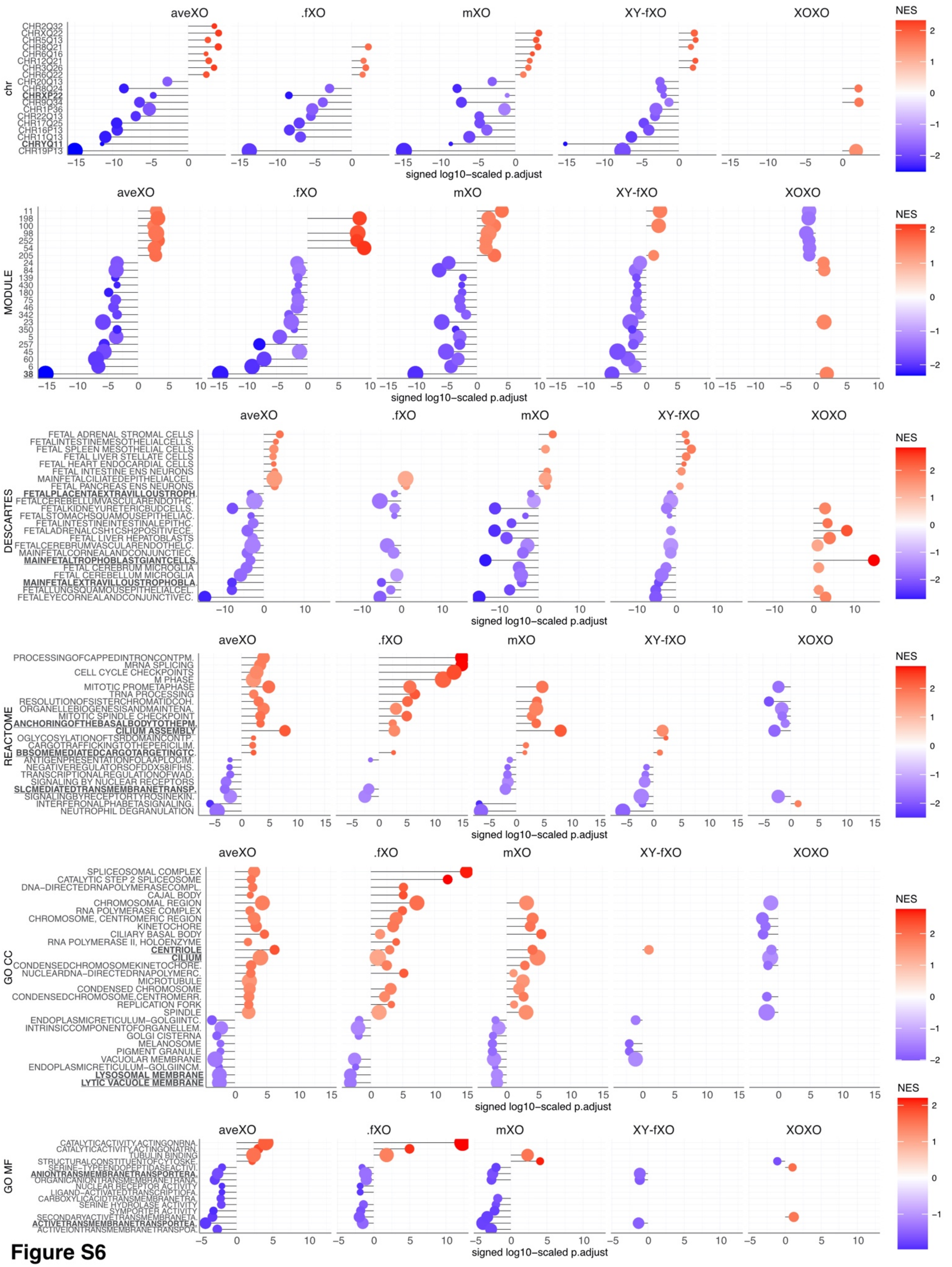
aveXO
.fXO
mXO
XY-fXO
XOXO
NES
chr
CHR2Q32
CHRXQ22
CHR5Q13
CHR8Q21
CHR6Q16
CHR12Q21
CHR3Q26
CHR6Q22
CHR20Q13
CHR8Q24
CHRXP22
CHR9Q34
CHR1P36
CHR22Q13
CHR17Q25
CHR16P13
CHR11Q13
CHRYQ11
CHR19P13
signed log10-scaled p.adjust
MODULE
DESCARTES
FETAL ADRENAL STROMAL CELLS
FETALINTESTINEMESOTHELIALCELLS.
FETAL SPLEEN MESOTHELIAL CELLS
FETAL LIVER STELLATE CELLS
FETAL HEART ENDOCARDIAL CELLS
FETAL INTESTINE ENS NEURONS
MAINFETALCILIATEDEPITHELIALCEL.
FETAL PANCREAS ENS NEURONS
FETALPLACENTAEXTRAVILLOUSTROPH.
FETALCEREBELLUMVASCULARENDOTHC.
FETALKIDNEYURETERICBUDCELLS.
FETALSTOMACHSQUAMOUSEPITHELIAC.
FETALINTESTINEINTESTINALEPITHC.
FETALADRENALCSH1CSH2POSITIVECE.
FETAL LIVER HEPATOBLASTS
FETALCEREBRUMVASCULARENDOTHELC.
MAINFETALCORNEALANDCONJUNCTIEC.
MAINFETALTROPHOBLASTGIANTCELLS.
FETAL CEREBRUM MICROGLIA
FETAL CEREBELLUM MICROGLIA
MAINFETALEXTRAVILLOUSTROPHOBLA.
FETALLUNGSQUAMOUSEPITHELIALCEL.
FETALEYECORNEALANDCONJUNCTIVEC.
REACTOME
PROCESSINGOFCAPPEDINTRONCONTPM.
MRNA SPLICING
CELL CYCLE CHECKPOINTS
M PHASE
MITOTIC PROMETAPHASE
TRNA PROCESSING
RESOLUTIONOFSISTERCHROMATIDCOH.
ORGANELLEBIOGENESISANDMAINTENA.
MITOTIC SPINDLE CHECKPOINT
ANCHORINGOFTHEBASALBODYTOTHEPM.
CILIUM ASSEMBLY
OGLYCOSYLATIONOFTSRDOMAINCONTP.
CARGOTRAFFICKINGTOTHEPERICILIM.
BBSOMEMEDIATEDCARGOTARGETINGTC.
ANTIGENPRESENTATIONFOLAAPLOCIM.
NEGATIVEREGULATORSOFDDX58IFIHS.
TRANSCRIPTIONALREGULATIONOFWAD.
SIGNALING BY NUCLEAR RECEPTORS
SLCMEDIATEDTRANSMEMBRANETRANSP.
SIGNALINGBYRECEPTORTYROSINEKIN.
INTERFERONALPHABETASIGNALING.
NEUTROPHIL DEGRANULATION
GO CC
SPLICEOSOMAL COMPLEX
CATALYTIC STEP 2 SPLICEOSOME
DNA−DIRECTEDRNAPOLYMERASECOMPL.
CAJAL BODY
CHROMOSOMAL REGION
RNA POLYMERASE COMPLEX
CHROMOSOME, CENTROMERIC REGION
KINETOCHORE
CILIARY BASAL BODY
RNA POLYMERASE II, HOLOENZYME
CENTRIOLE
CILIUM
CONDENSEDCHROMOSOMEKINETOCHORE.
NUCLEARDNA−DIRECTEDRNAPOLYMERC.
MICROTUBULE
CONDENSED CHROMOSOME
CONDENSEDCHROMOSOME,CENTROMERR.
REPLICATION FORK
SPINDLE
ENDOPLASMICRETICULUM−GOLGIINTC.
INTRINSICCOMPONENTOFORGANELLEM.
GOLGI CISTERNA
MELANOSOME
PIGMENT GRANULE
VACUOLAR MEMBRANE
ENDOPLASMICRETICULUM−GOLGIINCM.
LYSOSOMAL MEMBRANE
LYTIC VACUOLE MEMBRANE
GO MF
CATALYTICACTIVITYACTINGONRNA.
CATALYTICACTIVITYACTINGONATRN.
TUBULIN BINDING
STRUCTURALCONSTITUENTOFCYTOSKE.
SERINE−TYPEENDOPEPTIDASEACTIVI.
ANIONTRANSMEMBRANETRANSPORTERA.
ORGANICANIONTRANSMEMBRANETRANA.
NUCLEAR RECEPTOR ACTIVITY
LIGAND−ACTIVATEDTRANSCRIPTIOFA.
CARBOXYLICACIDTRANSMEMBRANETRA.
SERINE HYDROLASE ACTIVITY
SYMPORTER ACTIVITY
SECONDARYACTIVETRANSMEMBRANETA.
ACTIVETRANSMEMBRANETRANSPORTEA.
ACTIVEIONTRANSMEMBRANETRANSPOA.

**Figure S6**

**Figure S6:** Gene-set enrichment analysis (GSEA) against various MSigDB gene sets (in order: cytogenetic bands, computationally-derived modules, Descartes' human fetal cell type markers (6), Reactome and GO cellular component and molecular function). Five comparisons were included: fXO, mXO, XY-fXO, a gene list re-ranked by the average Wald statistic from all three quantile-normalized sets ("aveXO"), and a control comparison between male- and female-derived 45,X samples ("XOXO"). Bubble position, color and size, denote the signed log10-scaled GSEA p.adjust value, the normalized enrichment score, and the number of core genes driving the enrichment, respectively, and are plotted opposite of the enriched gene set titles (abbreviated).

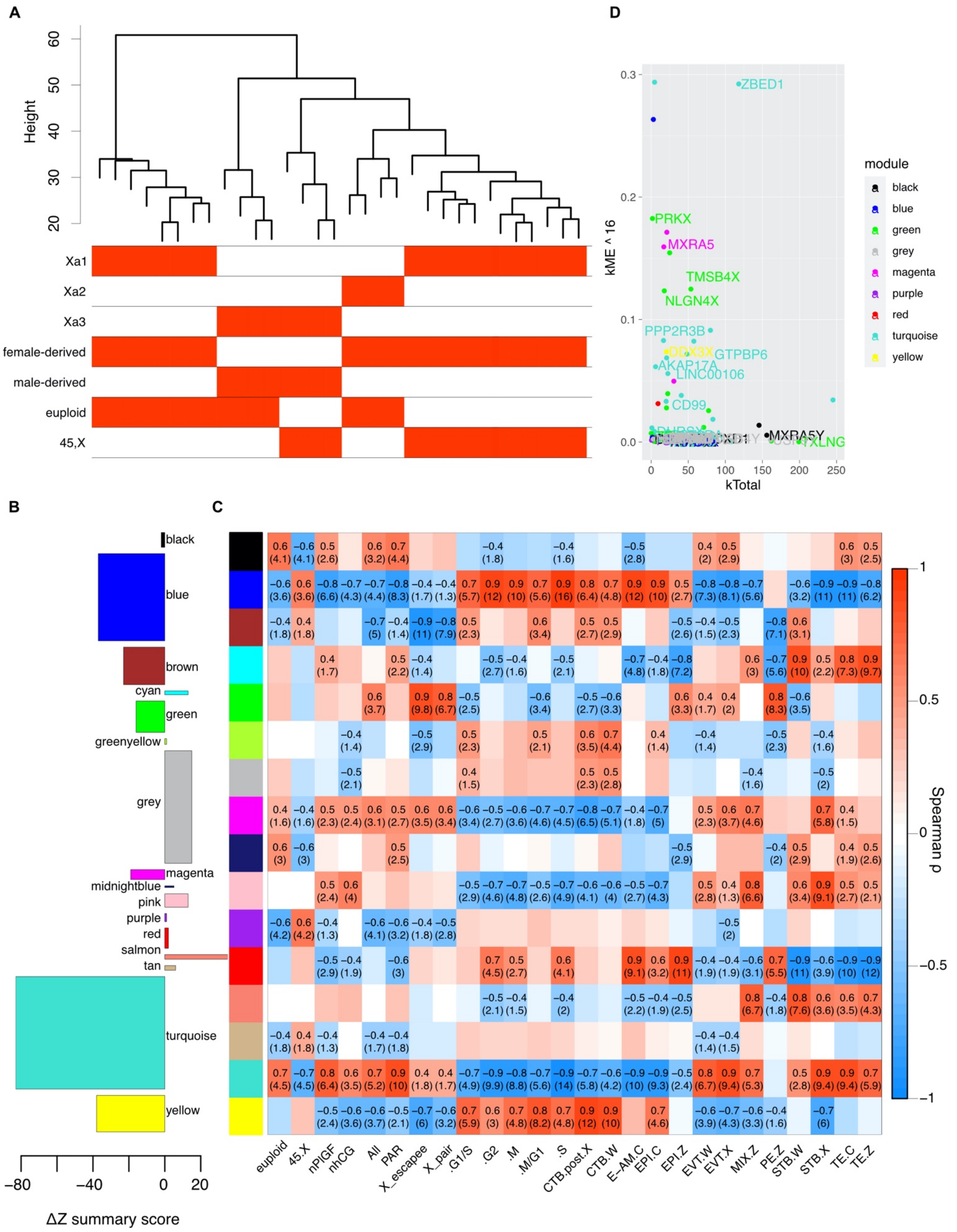
A
Height
60
50
40
30
20
Xa1
Xa2
Xa3
female-derived
male-derived
euploid
45,X
D
kME ^ 16
kTotal
0.3
0.2
0.1
0.0
0
50
100
150
200
250
ZBED1
PRKX
MXRA5
TMSB4X
NLGN4X
PPP2R3B
DDX3X
GTPBP6
AKAP17A
LINC00106
CD99
MXRA5Y
NLGN4Y
module
black
blue
green
grey
magenta
purple
red
turquoise
yellow
B
black
blue
brown
cyan
green
greenyellow
grey
magenta
midnightblue
pink
purple
red
salmon
tan
turquoise
yellow
−80
−40
0
20
ΔZ summary score
C
euploid
45,X
nPlGF
nhCG
All
PAR
X_escapee
X_pair
G1/S
G2
M
M/G1
S
CTB.post.X
CTB.W
E-AM.C
EPI.C
EPI.Z
EVT.W
EVT.X
MIX.Z
PE.Z
STB.W
STB.X
TE.C
TE.Z
Spearman ρ
1
0.5
0
−0.5
−1

**Figure S7**

**Figure S7: A.)** Clustering of all BAP samples for WGCNA separates euploid 46,XX lines with by karyotype before donor, except for XX6 samples reflecting the opposite XCI choice (Xa2). **B.)** Modules resolved in the full network (32 samples) are preserved significantly better (Z summary statistic) in a smaller subset (16 samples) reflecting mixed euploid and 45,X samples, than in a subset consisting exclusively of 45,X samples. **C.)** Correlation matrix (Spearman $\rho$) between WGCNA modules and categorical (euploid, 45,X), measured (PlGF, hCG) or aggregated cell type expression traits. The latter cover cell type markers from early human embryonic studies (2, 3, 7, 8), suffix-labeled by first initial (.W, .X, .Z, .C). Cell fates abbreviated for cytotrophoblast (CTB), early amnion (E-AM), epiblast (EPI), extravillous TB (EVT), mixed (MIX), primordial endoderm (PE) and syncytiotrophoblast (STB), as in Fig. 5. Significance estimate of each correlation denoted underneath $\rho$ in parentheses (as -log10(p), except where $p > 0.05$). **D.)** Correlation coefficient (kME, raised to network power of 16) for each PAR, PAIR or X-specific escapee with its assigned module's eigengene, plotted over its total connectivity (kTotal). Genes are colored to by assigned module, and labeled by gene name if homolog is present on Y. The top X/Y-linked hub gene for the turquoise module is PAR gene *ZBED1* (raw kME correlation of ~0.92).

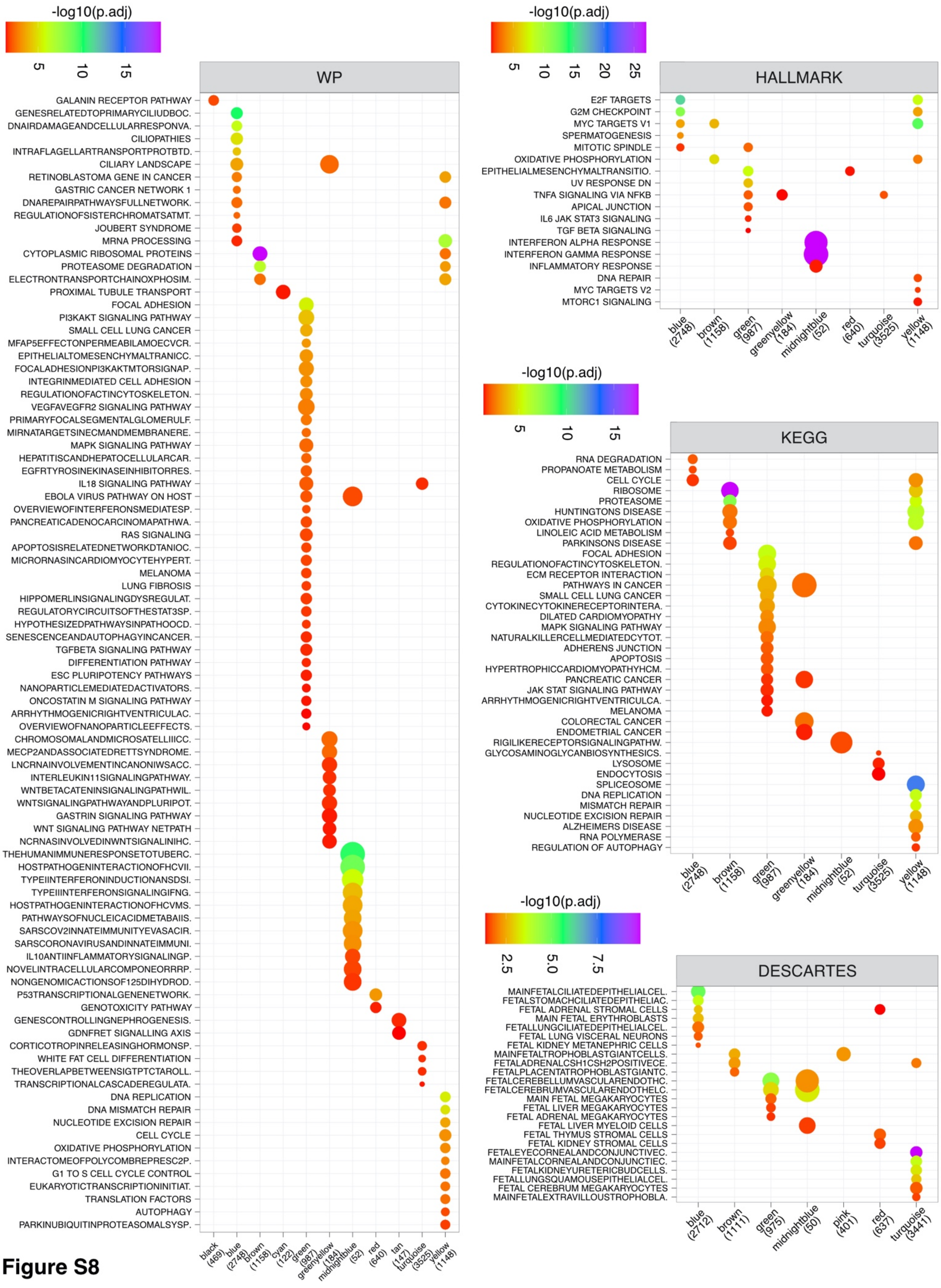
-log10(p.adj)
5
10
15
WP
GALANIN RECEPTOR PATHWAY
GENESRELATEDTOPRIMARYCILIUDBOC.
DNAIRDAMAGEANDCELLULARRESPONVA.
CILIOPATHIES
INTRAFLAGELLARTRANSPORTPROTBTD.
CILIARY LANDSCAPE
RETINOBLASTOMA GENE IN CANCER
GASTRIC CANCER NETWORK 1
DNAREPAIRPATHWAYSFULLNETWORK.
REGULATIONOFSISTERCHROMATSATMT.
JOUBERT SYNDROME
MRNA PROCESSING
CYTOPLASMIC RIBOSOMAL PROTEINS
PROTEASOME DEGRADATION
ELECTRONTRANSPORTCHAINOXPHOSIM.
PROXIMAL TUBULE TRANSPORT
FOCAL ADHESION
PI3KAKT SIGNALING PATHWAY
SMALL CELL LUNG CANCER
MFAP5EFFECTONPERMEABILAMOECVCR.
EPITHELIALTOMESENCHYMALTRANICC.
FOCALADHESIONPI3KAKTMTORSIGNAP.
INTEGRINMEDIATED CELL ADHESION
REGULATIONOFACTINCYTOSKELETON.
VEGFAVEGFR2 SIGNALING PATHWAY
PRIMARYFOCALSEGMENTALGLOMERULF.
MIRNATARGETSINECMANDMEMBRANERE.
MAPK SIGNALING PATHWAY
HEPATITISCANDHEPATOCELLULARCAR.
EGFRTYROSINEKINASEINHIBITORRES.
IL18 SIGNALING PATHWAY
EBOLA VIRUS PATHWAY ON HOST
OVERVIEWOFINTERFERONSMEDIATESP.
PANCREATICADENOCARCINOMAPATHWA.
RAS SIGNALING
APOPTOSISRELATEDNETWORKDTANIOC.
MICRORNASINCARDIOMYOCYTEHYPERT.
MELANOMA
LUNG FIBROSIS
HIPPOMERLINSIGNALINGDYSREGULAT.
REGULATORYCIRCUITSOFTHESTAT3SP.
HYPOTHESIZEDPATHWAYSINPATHOOCD.
SENESCENCEANDAUTOPHAGYINCANCER.
TGFBETA SIGNALING PATHWAY
DIFFERENTIATION PATHWAY
ESC PLURIPOTENCY PATHWAYS
NANOPARTICLEMEDIATEDACTIVATORS.
ONCOSTATIN M SIGNALING PATHWAY
ARRHYTHMOGENICRIGHTVENTRICULAC.
OVERVIEWOFNANOPARTICLEEFFECTS.
CHROMOSOMALANDMICROSATELLIIICC.
MECP2ANDASSOCIATEDRETTSYNDROME.
LNCRNAINVOLVEMENTINCANONIWSACC.
INTERLEUKIN11SIGNALINGPATHWAY.
WNTBETACATENINSIGNALINGPATHWIL.
WNTSIGNALINGPATHWAYANDPLURIPOT.
GASTRIN SIGNALING PATHWAY
WNT SIGNALING PATHWAY NETPATH
NCRNASINVOLVEDINWNTSIGNALINIHC.
THEHUMANIMMUNERESPONSETOTUBERC.
HOSTPATHOGENINTERACTIONOFHCVII.
TYPEIINTERFERONINDUCTIONANSDSI.
TYPEIIINTERFERONSIGNALINGIFNG.
HOSTPATHOGENINTERACTIONOFHCVMS.
PATHWAYSOFNUCLEICACIDMETABAIIS.
SARSCOV2INNATEIMMUNITYEVASACIR.
SARSCORONAVIRUSANDINNATEIMMUNI.
IL10ANTIINFLAMMATORYSIGNALINGP.
NOVELINTRACELLULARCOMPONEORRRP.
NONGENOMICACTIONSOF125DIHYDROD.
P53TRANSCRIPTIONALGENENETWORK.
GENOTOXICITY PATHWAY
GENESCONTROLLINGNEPHROGENESIS.
GDNFRET SIGNALLING AXIS
CORTICOTROPINRELEASINGHORMONSP.
WHITE FAT CELL DIFFERENTIATION
THEOVERLAPBETWEENSIGTPTCTAROLL.
TRANSCRIPTIONALCASCADEREGULATA.
DNA REPLICATION
DNA MISMATCH REPAIR
NUCLEOTIDE EXCISION REPAIR
CELL CYCLE
OXIDATIVE PHOSPHORYLATION
INTERACTOMEOFPOLYCOMBREPRESC2P.
G1 TO S CELL CYCLE CONTROL
EUKARYOTICTRANSCRIPTIONINITIAT.
TRANSLATION FACTORS
AUTOPHAGY
PARKINUBIQUITINPROTEASOMALSYSP.
black (469)
blue (2748)
brown (1158)
cyan (122)
green (987)
greenyellow (184)
midnightblue (52)
red (640)
tan (147)
turquoise (3525)
yellow (1148)
-log10(p.adj)
5
10
15
20
25
HALLMARK
E2F TARGETS
G2M CHECKPOINT
MYC TARGETS V1
SPERMATOGENESIS
MITOTIC SPINDLE
OXIDATIVE PHOSPHORYLATION
EPITHELIALMESENCHYMALTRANSITIO.
UV RESPONSE DN
TNFA SIGNALING VIA NFKB
APICAL JUNCTION
IL6 JAK STAT3 SIGNALING
TGF BETA SIGNALING
INTERFERON ALPHA RESPONSE
INTERFERON GAMMA RESPONSE
INFLAMMATORY RESPONSE
DNA REPAIR
MYC TARGETS V2
MTORC1 SIGNALING
blue (2748)
brown (1158)
green (987)
greenyellow (184)
midnightblue (52)
red (640)
turquoise (3525)
yellow (1148)
-log10(p.adj)
5
10
15
KEGG
RNA DEGRADATION
PROPANOATE METABOLISM
CELL CYCLE
RIBOSOME
PROTEASOME
HUNTINGTONS DISEASE
OXIDATIVE PHOSPHORYLATION
LINOLEIC ACID METABOLISM
PARKINSONS DISEASE
FOCAL ADHESION
REGULATIONOFACTINCYTOSKELETON.
ECM RECEPTOR INTERACTION
PATHWAYS IN CANCER
SMALL CELL LUNG CANCER
CYTOKINECYTOKINERECEPTORINTERA.
DILATED CARDIOMYOPATHY
MAPK SIGNALING PATHWAY
NATURALKILLERCELLMEDIATEDCYTOT.
ADHERENS JUNCTION
APOPTOSIS
HYPERTROPHICCARDIOMYOPATHYHCM.
PANCREATIC CANCER
JAK STAT SIGNALING PATHWAY
ARRHYTHMOGENICRIGHTVENTRICULCA.
MELANOMA
COLORECTAL CANCER
ENDOMETRIAL CANCER
RIGILIKERECEPTORSIGNALINGPATHW.
GLYCOSAMINOGLYCANBIOSYNTHESICS.
LYSOSOME
ENDOCYTOSIS
SPLICEOSOME
DNA REPLICATION
MISMATCH REPAIR
NUCLEOTIDE EXCISION REPAIR
ALZHEIMERS DISEASE
RNA POLYMERASE
REGULATION OF AUTOPHAGY
blue (2748)
brown (1158)
green (987)
greenyellow (184)
midnightblue (52)
turquoise (3525)
yellow (1148)
-log10(p.adj)
2.5
5.0
7.5
DESCARTES
MAINFETALCILIATEDEPITHELIALCEL.
FETALSTOMACHCILIATEDEPITHELIAC.
FETAL ADRENAL STROMAL CELLS
MAIN FETAL ERYTHROBLASTS
FETALLUNGCILIATEDEPITHELIALCEL.
FETAL LUNG VISCERAL NEURONS
FETAL KIDNEY METANEPHRIC CELLS
MAINFETALTROPHOBLASTGIANTCELLS.
FETALADRENALCSH1CSH2POSITIVECE.
FETALPLACENTATROPHOBLASTGIANTC.
FETALCEREBELLUMVASCULARENDOTHC.
FETALCEREBRUMVASCULARENDOTHELC.
MAIN FETAL MEGAKARYOCYTES
FETAL LIVER MEGAKARYOCYTES
FETAL ADRENAL MEGAKARYOCYTES
FETAL LIVER MYELOID CELLS
FETAL THYMUS STROMAL CELLS
FETAL KIDNEY STROMAL CELLS
FETALEYECORNEALANDCONJUNCTIVEC.
MAINFETALCORNEALANDCONJUNCTIEC.
FETALKIDNEYURETERICBUDCELLS.
FETALLUNGSQUAMOUSEPITHELIALCEL.
FETAL CEREBRUM MEGAKARYOCYTES
MAINFETALEXTRAVILLOUSTROPHOBLA.
blue (2712)
brown (1111)
green (975)
midnightblue (50)
pink (401)
red (637)
turquoise (3441)

**Figure S8**

**Figure S8:** Over-representation analysis of each module against the Wikipathway, Hallmark, KEGG and Descartes' human fetal cell type marker collection (6), next to abbreviated gene set titles. Bubble size and color denote gene ratio, and log10-scaled p.adjust, respectively.
